# The Multilocus Multispecies Coalescent: A Flexible New Model of Gene Family Evolution

**DOI:** 10.1101/2020.05.07.081836

**Authors:** Qiuyi Li, Celine Scornavacca, Nicolas Galtier, Yao-Ban Chan

**Affiliations:** School of Mathematics and Statistics / Melbourne Integrative Genomics, The University of Melbourne, Melbourne, 3010, Australia; Institut des Sciences de l’Evolution, Université Montpellier, CNRS, IRD, EPHE, Montpellier, 34095, France

**Keywords:** Gene duplication, gene loss, horizontal gene transfer, incomplete lineage sorting, multispecies coalescent, hemiplasy, recombination

## Abstract

Incomplete lineage sorting (ILS), the interaction between coalescence and speciation, can generate incongruence between gene trees and species trees, as can gene duplication (D), transfer (T) and loss (L). These processes are usually modelled independently, but in reality, ILS can affect gene copy number polymorphism, i.e., interfere with DTL. This has been previously recognised, but not treated in a satisfactory way, mainly because DTL events are naturally modelled forward-in-time, while ILS is naturally modelled backwards-in-time with the coalescent. Here we consider the joint action of ILS and DTL on the gene tree/species tree problem in all its complexity. In particular, we show that the interaction between ILS and duplications/transfers (without losses) can result in patterns usually interpreted as resulting from gene loss, and that the realised rate of D, T and L becomes non-homogeneous in time when ILS is taken into account. We introduce algorithmic solutions to these problems. Our new model, the *multilocus multispecies coalescent* (MLMSC), which also accounts for any level of linkage between loci, generalises the multispecies coalescent model and offers a versatile, powerful framework for proper simulation and inference of gene family evolution.

---

Species trees and gene trees are two important kinds of phylogenetic trees which are key to the study of gene and genome evolution. A species tree depicts the evolutionary history of a set of organisms, whereas a gene tree depicts the evolutionary history of a gene family within a set of organisms. A gene tree is thus constrained by the species tree, which describes the evolution of the organisms containing the gene family.

When considering speciations as the only possible events shaping species histories (e.g., no hybridisation or reassortment possible), internal nodes of species trees represent only speciation events, and branch lengths represent divergence times or substitutions per site. Nodes of gene trees, in contrast, can be the result of a combination of diverse processes, such as variation in gene copy number or gene transfers between species, in addition to speciations. For this reason, gene trees often differ from species trees, which can be seen as a problem — if the goal is to infer the species tree — or a source of information — if the goal is to study molecular evolution.

These evolutionary processes can include several ‘gene-range’ events such as gene duplications, gene losses, and horizontal gene transfers. A gene duplication (D) is an event in which a single gene copy gives rise to two copies at distinct loci: the parent locus and a new (child) locus. In contrast, a gene loss (L) removes a gene from the genome. Horizontal gene transfer (T) occurs when a gene from one species enters the genome of another contemporary species, which can occur (frequently in bacteria, for example) through a number of biological mechanisms such as transformation, transduction, and conjugation. Collectively, we refer to these processes as ‘DTL’. These events can occur multiple times, allowing the gene tree to possibly differ greatly from the species tree. Examples of these events are given in Figure 1.

**Figure 1:**
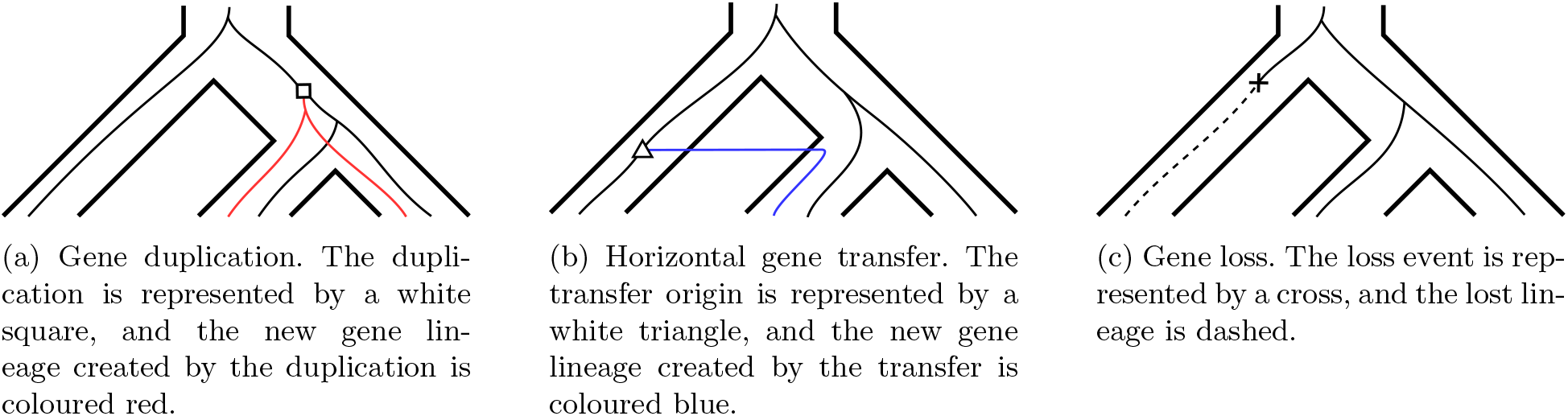
Tree representations of a gene duplication, a horizontal gene transfer, and a gene loss, respectively. The gene lineages (finer lines) evolve within a species tree (outer ‘tubes’).

In addition to these events, a gene tree can also be incongruent to the species tree due to a phenomenon known as incomplete lineage sorting (ILS, Maddison, 1997). When a population of individuals undergoes several speciations in a relatively short time, polymorphism (different alleles) maintained throughout this time may eventually fix in different descendant lineages. This can produce discrepancies between the gene tree and species tree. ILS is more likely to occur in branches of the species tree (i.e., ancestral species) that represent small time spans and/or large population sizes (Pamilo and Nei, 1988). An example of ILS is given in Figure 2. Hemiplasy (Avise and Robinson, 2008) is a term used to refer to the species tree/gene tree conflicts that result from incomplete lineage sorting.

**Figure 2:**
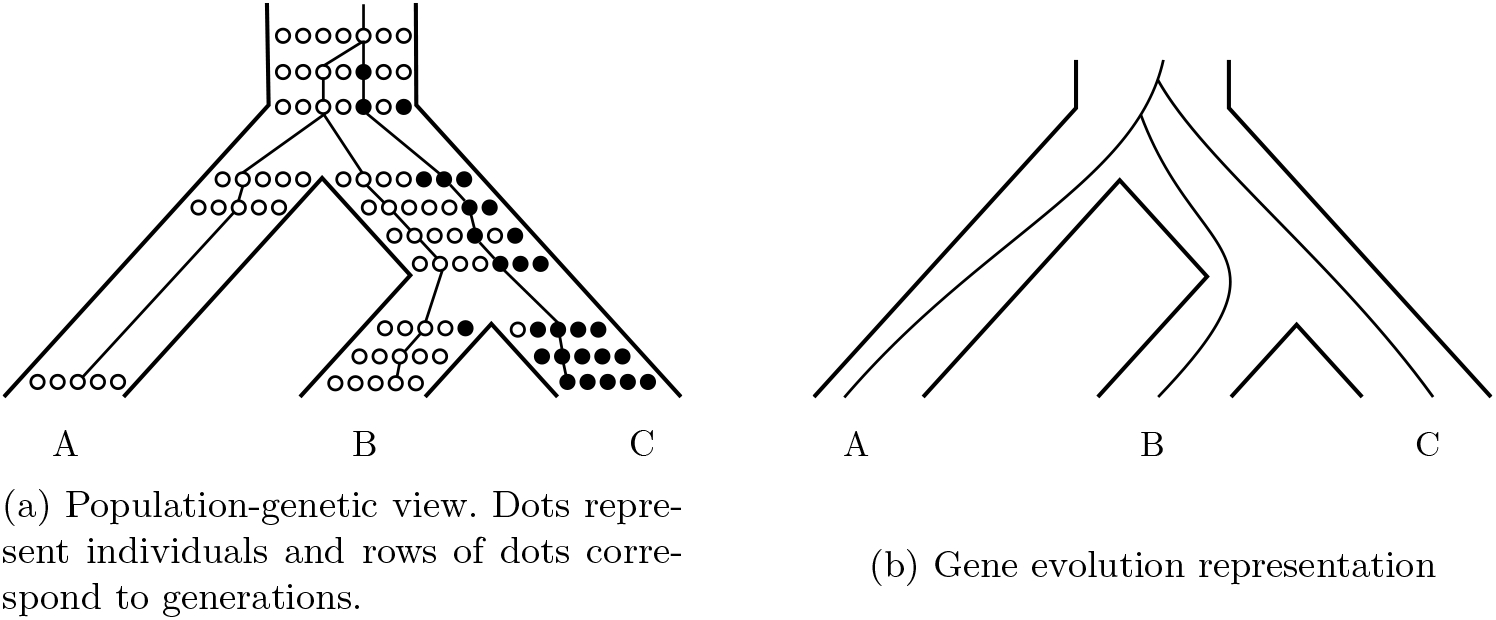
Example of incomplete lineage sorting (ILS). The original population contains a single white allele for the gene of interest. First, a mutation leads to a new black allele at the locus, then the first speciation takes place, rapidly followed by a second one. As the white and black alleles still coexist when the second speciation takes place, both alleles may be fixed in separate descendant species, resulting in a gene history which differs from the species history.

Coalescent theory (Kingman, 1982) provides a genealogical interpretation of ILS that helps to connect this phenomenon to gene tree-species tree discordance. A key point is that the age of the common ancestor of two gene copies sampled in two different species is older (in the absence of DTL) than the time of speciation between the two species. This is due to the existence of polymorphism in the ancestral species. When speciations occur far apart in time from each other, ancestral polymorphism is unlikely to result in differences in the topology of gene trees and the species tree, although it will create differences in branch lengths. If, however, two or more speciations occur in a time interval of the order of coalescence times, then coalescence and speciation may interact. This can cause not only branch lengths, but also topology, to differ between gene trees and species trees, as in Figure 2.

The multispecies coalescent model (MSC, Rannala and Yang, 2003) predicts the effect of ILS on gene tree branch lengths and topology as a function of the effective population size and timing of speciations. ILS and the MSC have received much attention after the discovery that classical phylogenetic methods are inconsistent in a subset of the parameter space (Kubatko and Degnan, 2007; Roch and Steel, 2015), and in the context of the study of convergent evolution (Guerrero and Hahn, 2018).

So far, ILS has mainly been considered separately from other sources of conflict between gene and species trees. However, ILS can interact in complex and often unintuitive ways with the processes of gene duplication, loss, and transfer, as first noted by Rasmussen and Kellis (2012). This is because DTL events spend some time in a polymorphic stage in a population, when individuals differ from each other in terms of gene copy number, before they become fixed. If speciations occur during this transient period of polymorphism, then the issue of lineage sorting becomes even more complex than in the one-locus case. For example, an allele which is not yet fixed can be lost, as shown in Figure 3. It is also possible that a newly created locus does not fix in all descendant species, as shown in Figure 4. Thus modelling ILS together with DTL requires greater flexibility than simply modelling each process individually. We refer to the discrepancies in gene copy number that result from the interaction between ILS and DTL as ‘copy number hemiplasy’ (or CNH for short). We note here that Rasmussen and Kellis’ model considers DTL and ILS independently, and does not model hemiplasy. Therefore, it cannot handle the scenarios shown in Figures 3 and 4.

**Figure 3:**
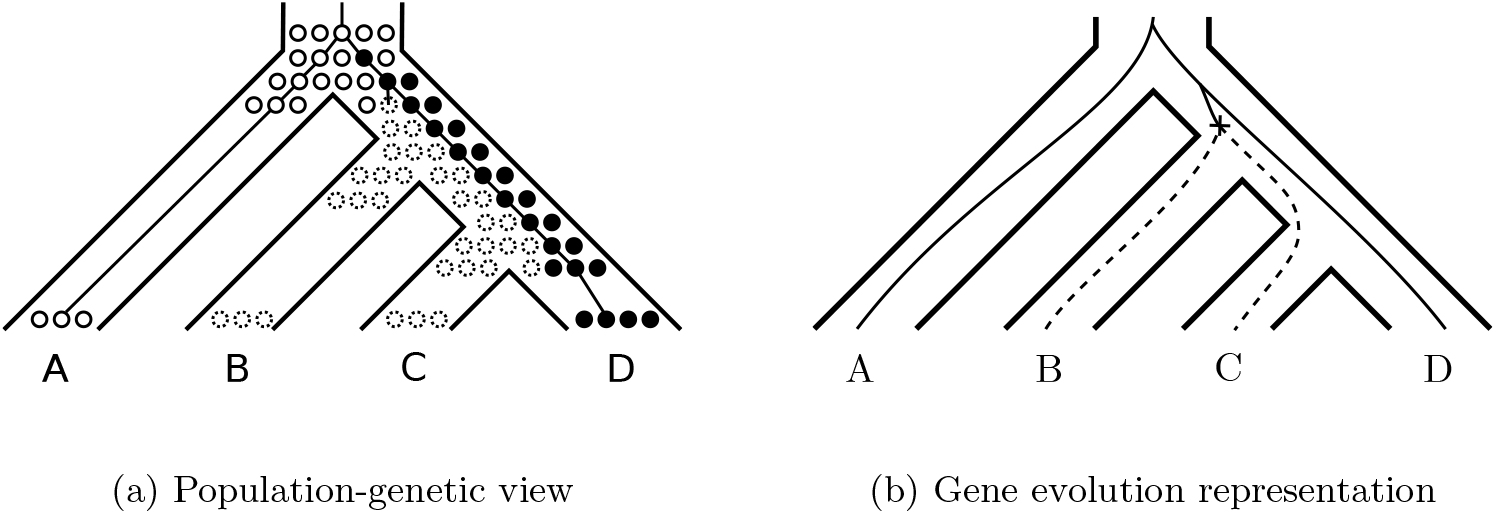
Example of ILS interacting with loss. With only one loss event, two descendant species may end up with an empty locus (represented by dotted circles) due to the presence of ILS.

**Figure 4:**
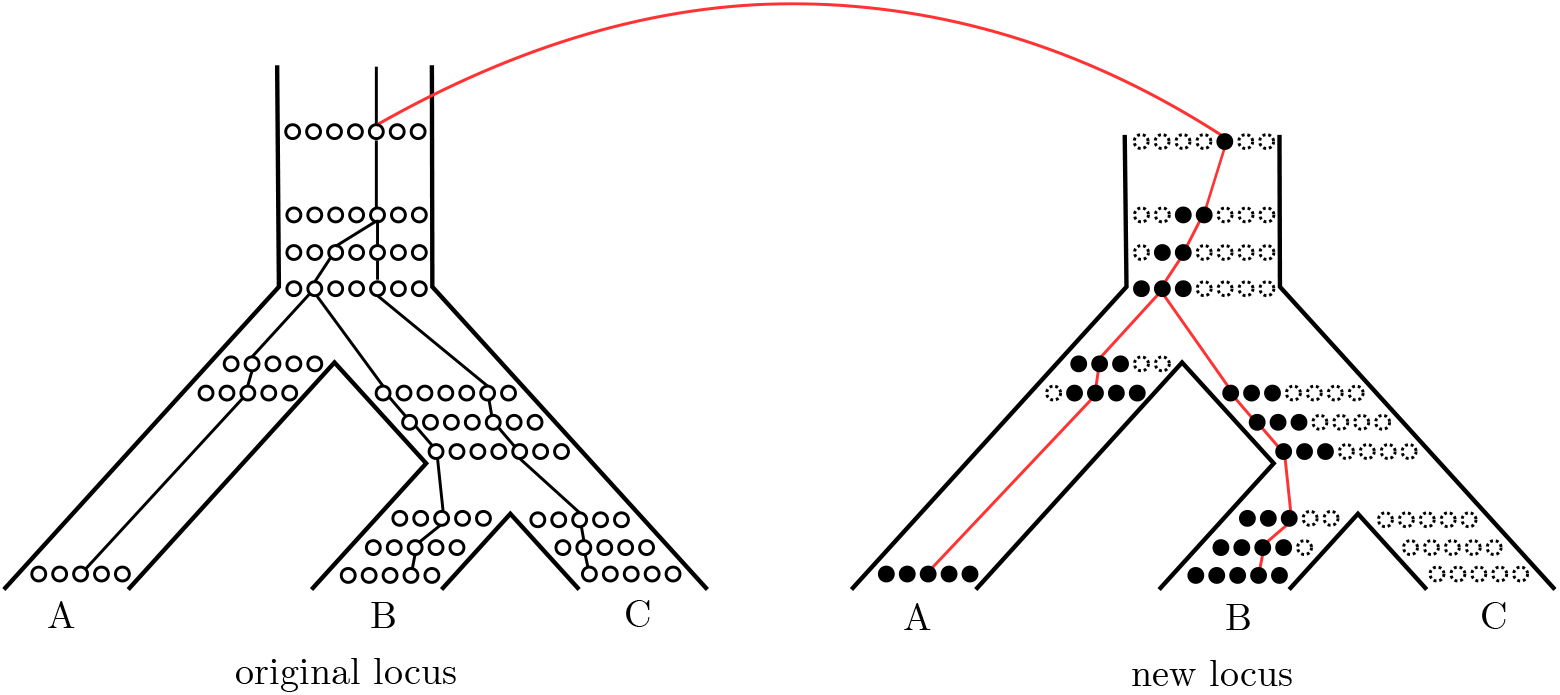
Example of copy number hemiplasy. A duplication arises in an ancestral species, and then two successive speciation events occur. This results in some species retaining two copies of the gene, while others have only one, without any gene loss event. Here, the original gene descends to all extant species (*A, B*, and *C*) in the original locus, but the duplicated gene only fixes in species *A* and *B*. Species *B* ends up with the same number of gene copies as *A*, while being more closely related to *C*.

Discrepancies between a gene tree and a species tree due to DTL and/or ILS are frequently analysed via mappings from the gene tree into the species tree, called *reconciliations* (Goodman et al., 1979). More formally, a reconciliation between a gene tree *G* and species tree *S* is a mapping of the nodes of *G* into the nodes (or a sequence of nodes) of *S* (Doyon et al., 2011), respecting some constraints that depend on the evolutionary model chosen. This gives rise to the problem of reconciliation inference, where we seek to reconstruct the ‘true’ reconciliation from the gene and species trees.

There are two main paradigms for reconciliation inference: parsimonious and probabilistic (Doyon et al., 2011). In the parsimonious approach, we assign a cost to each evolutionary event, and search for a most parsimonious reconciliation, i.e., one which induces the lowest total cost. In the probabilistic approach, a stochastic model of evolution is assumed, and either the reconciliation with the maximum likelihood under this model is found, or a Bayesian approach is used to sample the posterior reconciliation space. In general, the probabilistic approach is more accurate but less time efficient, since it typically requires the estimation of many parameters, e.g., effective population sizes and branch lengths.

In our discussion, it is important to distinguish between three concepts:

- The *model* of evolution — a specification of what can (and cannot) happen in the evolution of a gene family;
- A specification of the statistical processes involved in producing the effects captured by the model, which we call here the *generating process* of the model, which is necessary to support a probabilistic method; and
- A *method* to choose an optimal reconciliation under some criterion (parsimonious or probabilistic).

Every reconciliation method assumes a particular model of gene family evolution, limiting the potential sources of gene tree-species tree discrepancy. However, the underlying model is not always explicitly specified, which is necessary for a proper understanding and comparison of reconciliation methods.

The first reconciliation models considered only gene duplications and losses as the sources of discrepancy (Goodman et al., 1979; Zhang, 1997; Arvestad et al., 2004; Durand et al., 2006; Rasmussen and Kellis, 2011). More recently, horizontal gene transfer has been included in these models (Arvestad et al., 2009; Doyon et al., 2010; David and Alm, 2011; Tofigh et al., 2011; Sjöstrand et al., 2013), increasing both their complexity and realism. Statistically, these models are usually generated by birth-death processes running inside the species tree. On the other hand, the multispecies coalescent (Rannala and Yang, 2003) generates a model where discrepancies between gene and species trees are only due to ILS.

Only a few models (and corresponding methods to infer reconciliations) exist which attempt to unify these processes. The DLCoalRecon and DLCpar methods (Rasmussen and Kellis, 2012; Wu et al., 2014; Du et al., 2019; Mawhorter et al., 2019) consider ILS together with duplications and losses, overlaid on a model called DLCoal. SimPhy (Mallo et al., 2015) is a simulator based on the DLCoal model that additionally considers transfers. Schrempf and Szöllősi (2018) also consider these possible events, but use a different model that we call here the *haplotype tree model*. Lastly, the IDTL (Chan et al., 2017) and Notung (Stolzer et al., 2012) methods incorporate both ILS and transfers with duplications and losses, but again using different underlying models. Each of these models are slightly different from each other, and some of the papers (Chan et al., 2017; Du et al., 2019) compare the various models. However, they all have their limitations, which are discussed in Section “Existing Models of Evolution”. In particular, none of them can appropriately model copy number hemiplasy, although CNH is at the heart of the ILS/DTL interaction.

When modelling ILS together with DTL, one must also consider the issue of genetic linkage between gene copies. A new gene copy arising from a duplication often appears close to the parent gene on the chromosome, in which case the two loci are expected to follow correlated coalescent processes. The strength of the correlation is controlled by the amount of recombination between the two loci; in the absence of recombination, the two genealogies will be the same. The joint coalescent process of partly linked loci in a single population is well characterised (Hudson, 1983), but the connection with speciation and ILS has only been rarely considered so far, the model of Slatkin and Pollack (2006) being one notable exception. (See the Supplementary Material for details about their model and how it relates to ours.) A realistic model of gene family evolution should account for the possible existence of linkage between duplicated gene copies.

In this paper, we propose a new gene family evolution model, called the *multilocus multispecies coalescent*, or MLMSC for short. (This should not be confused with the multilocus coalescent, or MLC for short, introduced by Rasmussen and Kellis (2012).) This model generalises the multispecies coalescent to gene families, and is designed to capture all possible scenarios that can arise through ILS, DTL, and interaction between these processes. The MLMSC combines forward- and backward-in-time modelling in order to properly account for copy number hemiplasy and linkage between loci. Importantly, we show that the realised rates of D, T and L become non-homogeneous as these processes interact with ILS, and we introduce a solution to this problem. The MLMSC model is more flexible and predicts a more diverse range of biological patterns than existing models of gene family evolution.

## Existing Models of Evolution

We consider the problem of modelling gene family evolution in a phylogenetic context. Given a set of species, we assume that we have sampled exactly one haploid genome per species, and these genomes contain all the gene copies descending from one particular gene present at the root of the species tree, i.e., a complete gene family. We aim to model the genealogical relationships between these gene copies, assuming that it has been shaped by speciations, DTL, and ILS. In this section, we first review the existing models which have previously addressed this problem.

### DLCoal Model

The reconciliation methods DLCoalRecon and DLCpar are both based on the DLCoal model (Rasmussen and Kellis, 2012). In this model, when a gene duplication occurs, the child and parent gene copies evolve independently of each other, i.e., the two loci are unlinked. Biologically, this arises when there is a sufficient level of recombination between the two loci that they can be considered to evolve completely independently.

To generate this model statistically, given a species tree, we first generate a tree under a duplication-loss birth-death process on the species tree. Beginning with the original species tree, at each duplication the tree is copied from that point onwards and attached to itself (where it may be subject to further duplications), and at each loss the tree is truncated. This produces the so-called *locus tree*, which depicts the bifurcating evolution of all loci containing a copy of the gene.

The gene tree is then constructed by applying a multispecies coalescent process within the locus tree, with the caveat that only one gene copy can ‘travel’ along the branch connecting a duplicated locus back to its parent. In other words, all the lineages in one particular locus must coalesce into one lineage before coalescing with a copy in its parent locus — the so-called *bounded coalescent process*. This does not permit copy number hemiplasy — a duplicated gene is transmitted to all descendant species at the new locus, barring gene loss. Under this model, ILS does not interfere with gene duplication and loss in generating variation in gene copy number among species.

An example of this model is given in Figure 5, in which a new locus (with descendants in *B* and *C*) is created by duplication. The duplicated gene is then fixed in all species at the new locus.

**Figure 5:**
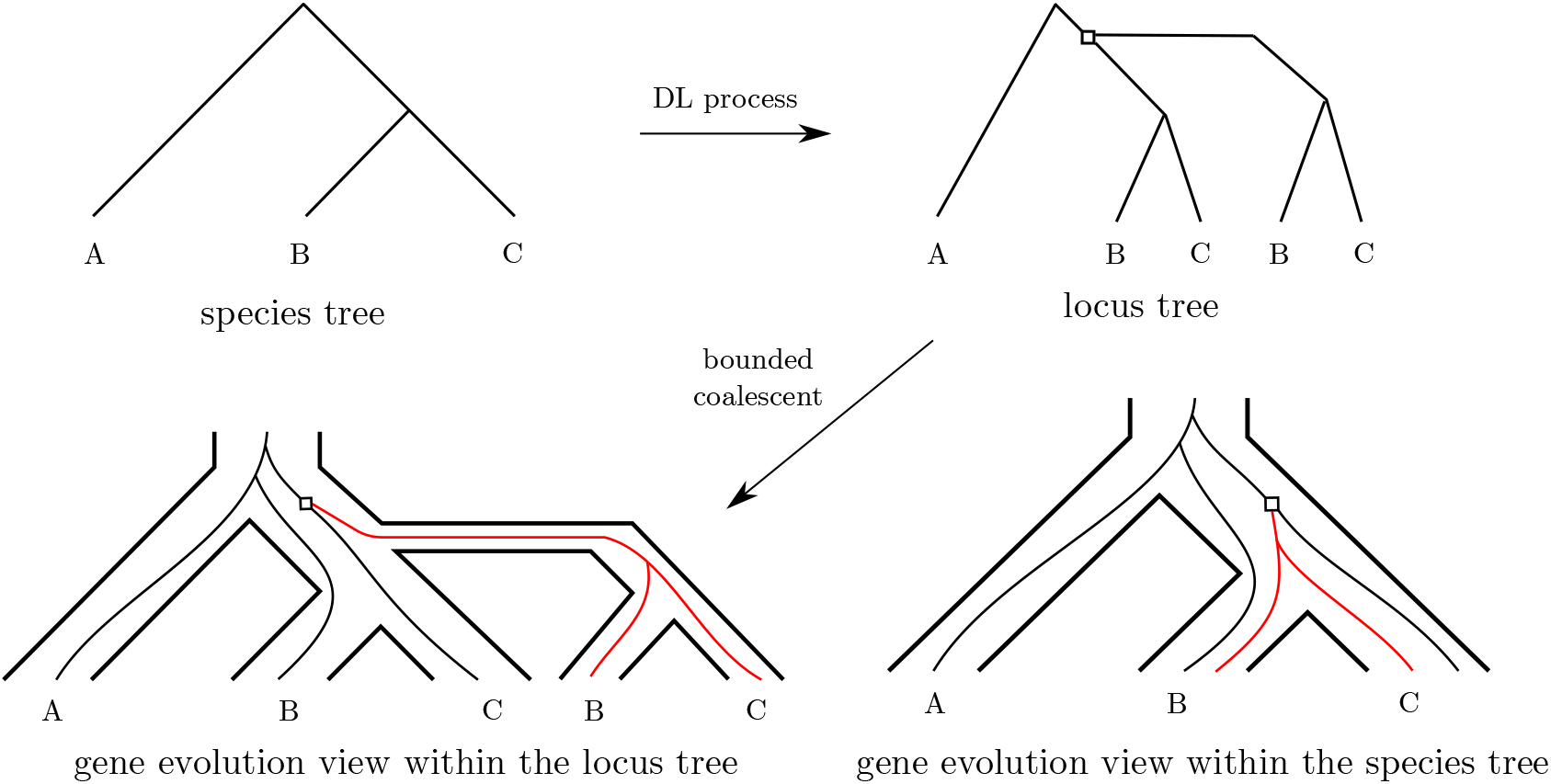
An example of the DLCoal model. Given a species tree, a locus tree is generated by applying a duplication-loss birth-death process on the species tree: the tree is copied from the point of duplication (white square) and attached to the original tree. The gene tree is then constructed under a bounded coalescent process within the locus tree, which requires all the lineages in the new locus (in red) to coalesce before entering the parent locus.

It is easy to extend this formulation to include gene transfers, and indeed SimPhy (Mallo et al., 2015) is a simulator that generates the DLCoal model, extended to transfers (note that SimPhy does not provide a method to find an optimal reconciliation).

### Haplotype Tree Model

The haplotype tree model was introduced by Schrempf in a talk at the SMBE 2018 conference (Schrempf and Szöllősi, 2018). In contrast to the DLCoal model, the haplotype tree model assumes that there is no recombination. This implies that a duplicated gene must undergo the exact same genealogical history as its parent gene — a strong assumption.

To generate this model statistically, we first generate a tree under the multispecies coalescent model on the species tree, obtaining a so-called *haplotype tree*. The gene tree is then obtained by performing a duplication-loss birth-death process on the haplotype tree. An example is given in Figure 6. Note that, while some sort of copy number hemiplasy is allowed in this model, it is restricted: a gene that duplicates must be sorted into exactly the same descendants as its parent gene. For instance, it is not allowed for a duplicated gene to be sorted into all descendant species if the parent gene is not.

**Figure 6:**
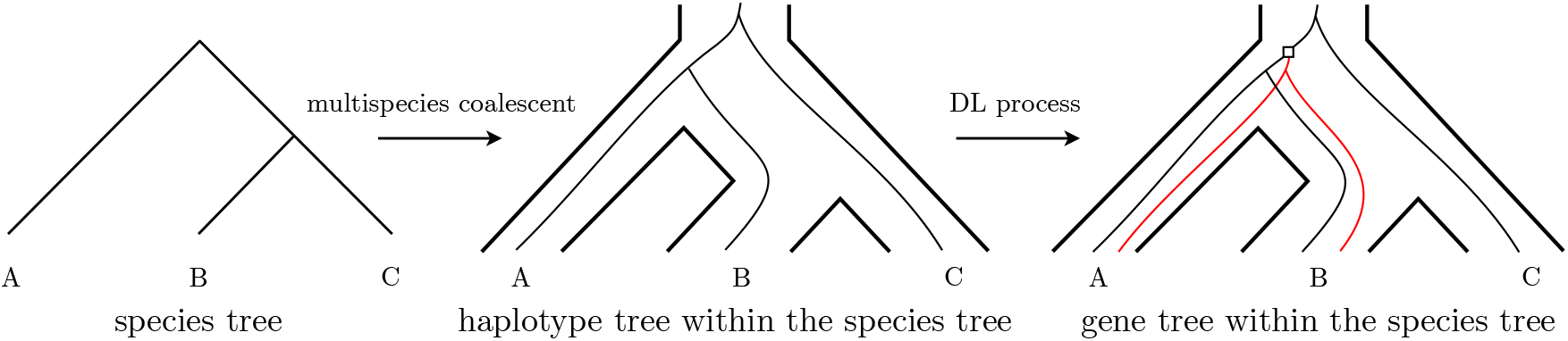
An example of the haplotype tree model. Given a species tree, a haplotype tree is constructed by applying a multispecies coalescent process within the species tree. The gene tree is then generated under a duplication-loss birth-death process on the haplotype tree: the haplotype tree is copied from the point of duplication (white square) and attached to the original tree. The new genes created by the duplication are coloured red.

### IDTL Model

The IDTL reconciliation method (Chan et al., 2017) takes a parsimony approach to the reconciliation problem. It defines events forward in time and assumes an underlying gene family evolution model which shares some elements with both the locus and haplotype tree models. Note that, while the model is specified, no generating process has been specified by Chan et al. (2017), since this is not necessary for a parsimony method.

One possible process based on the underlying model is to generate a tree under the multispecies coalescent model on the species tree, followed by a DTL birth-death process, as is done in the haplotype tree model. However, for each new locus, a new multispecies coalescent process is simulated as in the DLCoal model. As in the haplotype tree model, a limited form of copy number hemiplasy is allowed where a duplicate gene must be sorted into exactly the same descendant species as the parent gene. An example of this is given in Figure 7.

**Figure 7:**
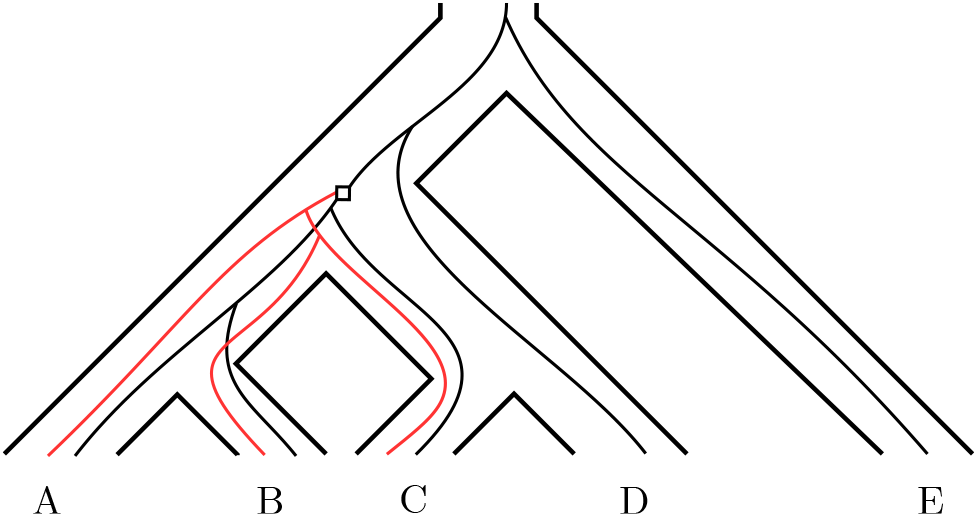
An example of the IDTL model. An allele which is sorted into species *A*, *B*, and *C* is duplicated. The duplicated gene must then be sorted into the same species, but the coalescent process may be different from the parent gene, resulting in a subtree (coloured red) with differing topology.

This model deals with recombination in an inconsistent way; this will be discussed further in Section “MLMSC vs IDTL Model”.

### Notung

The Notung method (Vernot et al., 2008; Stolzer et al., 2012) was one of the earliest methods to unify the processes of DTL and ILS. It is primarily intended to be used with non-binary species trees, i.e., situations where the branching order is unclear. Because of this, ILS in Notung is only allowed at a polytomy (node with more than two descending lineages) in the species tree, and each possible sorting of genes at the polytomy is considered to be equally likely. Thus there is no explicit modelling of alleles co-existing inside a species branch. As with the IDTL method, Notung is a parsimony method and does not have a formal generating process.

## The MLMSC Model

The models of gene family evolution in the literature do not model copy number hemiplasy or linked loci. Here we introduce a new gene family evolution model, the *multilocus multispecies coalescent* (MLMSC), for this purpose. We will specify a generating process for this model in the next section.

One of the primary conceptual difficulties in unifying ILS with a DTL model has been that ILS is traditionally generated by a backwards-in-time coalescent model, whereas the birth-death process used to generate DTL events runs forwards in time. The MLMSC instead uses a Wright-Fisher model for the population-level genealogical process, which also runs forwards in time and is therefore easier to merge with a birth-death process.

We first define some terminology: in the MLMSC model, one or more *species* exist, each of which are represented by a *population* consisting of a number of haploid *individuals*, or *members*. Each of these individuals carry one or more *loci*, which are fixed positions in the genome in which a *gene* may reside. We trace the evolution of a single *gene family*; that is, all descendants of a single gene, located in a locus inside an individual belonging to a single ancestral species. (For the purposes of the model, it is unimportant how this gene originated.) In order to describe the model, we describe the evolution of species, loci, and finally genes within populations in turn.

### The Evolution of Species

New species are created through *speciation*. When a speciation occurs, two new species are created, each of which is represented by a population which contains the same loci as the parent species. The two new species then continue evolving independently (as described below) as if they were continuations of the parent species, as in the MSC (Rannala and Yang, 2003).

### The Evolution of Loci

New loci are created through *gene duplication* (D) or *gene transfer* (T):

- A duplication occurs in a locus of an individual in a species. When it occurs, a new locus is created in that species. The new locus may be *linked* or *unlinked* to its parent locus (resulting from *linked duplication* or *unlinked duplication*, respectively); in the former case, it is also (indirectly) linked to all loci that its parent is linked to. The new locus is unlinked to all other existing loci.
- When a transfer occurs, the same process is followed as for a duplication, except that a new locus is instead created in a different but contemporary species of the originating individual, and this locus is unlinked to any existing locus.

A locus is lost in a species when there is no longer any individual of the species carrying a gene copy at that locus.

We can thus partition the set of all loci into *sets of linked loci*, which contain loci linked to each other but not to any locus outside the set. These sets can be of size 1, i.e., contain a single locus which is unlinked to all other loci. Note that loci within such a set are connected by a hierarchical relationship, with one locus (the one created first) being ancestral to the whole set (the *root locus* of the set), and each of the other loci descending from a parent locus within the set.

### The Evolution of Genes Within Populations

Within a population, genes are transmitted from generation to generation according to a Wright-Fisher model with recombination and gene loss. More precisely, for each locus, each individual in a generation inherits that locus from a parent individual in the previous generation. The parent individual is chosen independently for each set of linked loci. Within each set of linked loci, a parent individual is first randomly chosen for the root locus of the set. Then, recursively, each other locus may (with a certain probability) either descend from the same individual as its parent locus, or (representing a recombination between the two loci) another randomly chosen individual from the previous generation. An example of this process is given in Figure 8.

**Figure 8:**
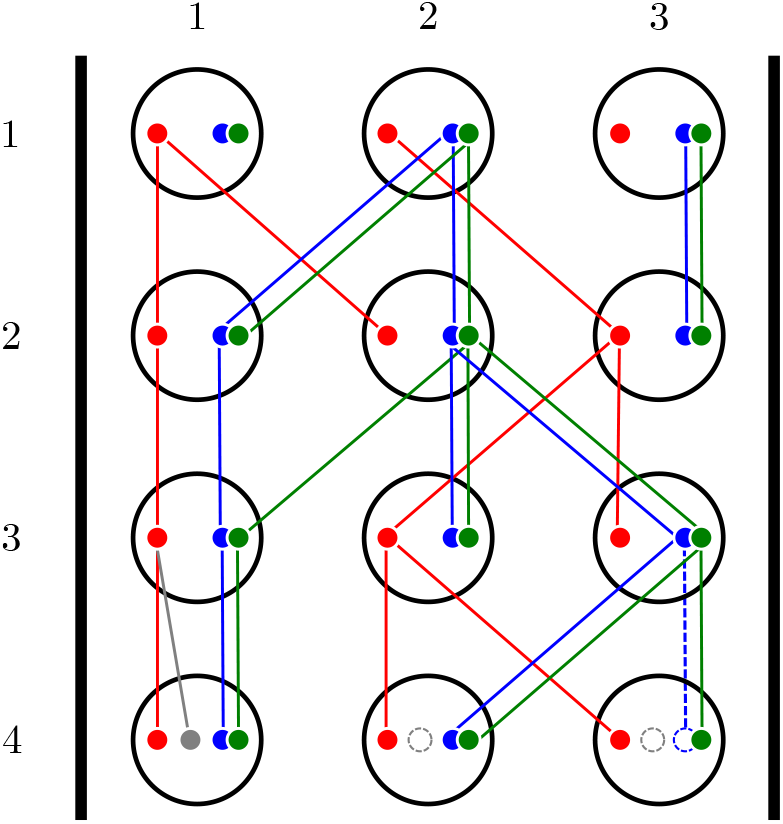
An example of the Wright-Fisher model with recombination and gene loss. In the first generation, each individual contains three loci: the blue and green loci are linked to each other and are inherited together, while the red locus is unlinked to the others and is inherited separately. At generation 3, a recombination event occurs between the blue and green loci, and individual 1 inherits these loci from different ancestors. At generation 4, an unlinked duplication occurs in individual 1, and a new locus (in grey) is created in all individuals in the population; however, only individual 1 contains a gene copy at that locus. A gene loss also occurs in the blue locus of individual 3.

Loci within a set of linked loci are therefore considered such that parent loci are treated before descendant loci. This order reflects the history of locus creation, and is not always interpretable in terms of the relative positions of loci along a linear chromosome. Any pair of linked loci thus evolves according to the two-locus Wright-Fisher model with recombination (Griffiths, 1991).

When an individual inherits a locus from an individual in the previous generation, with fixed probability the gene is lost, i.e., is not transmitted to the child individual (see Figure 8). When there is no longer any individual that carries a gene at a given locus in a population, that locus is lost from the species.

New genes are created through gene duplication and gene transfer, and passed through speciation:

- When a duplication or transfer occurs, a new locus is created in the population. The newly created gene is initially present only in a single individual in this locus (the individual undergoing duplication, or receiving the transfer).
- When a speciation occurs, the first generation of each of the child species descends independently from the last generation of the parent species as described above, and the process continues independently in the two newly created species.

## The Generating Process of the MLMSC Model

We now detail a statistical generating process of the MLMSC model under the following assumptions:

- A neutral Wright-Fisher model of evolution (no selection or segregation);
- Constant, large population size;
- Constant rates of duplication, transfer, loss, and recombination;
- Constant probability of a duplication being linked or unlinked.

Although the MLMSC model uses the Wright-Fisher model of evolution within a population, the coalescent (which approximates the Wright-Fisher model for *N ≫ n*) is used to generate the gene trees. Firstly, we introduce the concepts of *unilocus trees* and *haplotype trees and forests*.

*Unilocus trees model the history of duplications, transfers, and speciations.* For each locus, we define its *unilocus* tree as the subtree of the species tree rooted at the species where the locus first appears. The leaves of a unilocus tree thus correspond to the same locus in different species, and they are the only extant species which can inherit this locus. The history of gene duplications, gene transfers, and speciations in the MLMSC is thus stored as a collection of disconnected unilocus trees.

Note that this idea only slightly differs from the locus tree of the DLCoal model (see above): a collection of unilocus trees can equivalently be seen as a decorated locus tree, where the decorations are duplications/locus changes. For ease of understanding, in the following we consider them as disconnected trees.

*Haplotype trees and forests model the genealogies of lineages within a unilocus tree.* For a given locus, we model the genealogical relationships of gene copies across species via a collection of *haplotype trees*, each of which depicts the genealogy of the genes for a set of species. A *haplotype forest* is a set which contains either one haplotype tree or a number of disconnected haplotype trees, i.e., haplotype trees whose sets of leaves are disjoint. Generally, only one of the trees in the haplotype forest will actually describe the evolution of the gene, in which case we refer to it as *the* haplotype tree for the locus.

The haplotype forest for a locus is constrained by (evolves within) the unilocus tree for that locus. Haplotype trees and forests will be used to track the presence of different alleles in the populations (via ILS, CNH, and losses). Note that in our model, DTL events are modelled forwards in time and then haplotypes are modelled backwards in time.

To generate a gene tree from a species tree under the MLMSC model, we start with the species tree *S* as the unilocus tree for the original locus, and generate a haplotype tree for it. Then we recursively generate new events, new loci (with unilocus trees), and haplotype trees and forests for those loci, until all loci have been created. The haplotype trees are then concatenated together to form the full gene tree.

### Generating a Haplotype Tree for the Original Locus

In the original locus, the unilocus tree is the original species tree *S*. For this locus only, the haplotype tree is generated according to the standard multispecies coalescent, starting from a single copy of the gene in each leaf of the tree. We also set the haplotype forest to be the set containing only the haplotype tree.

For loci created by duplication or transfer, a more complex process is required. We first describe how to generate new loci and unilocus trees, then return to generating haplotype trees within those unilocus trees.

### Simulating Surviving Events

After we have generated a haplotype tree and forest for a locus, we simulate DTL events which occur in that locus. It is important to realise that we do not want to simulate all DTL events that occur, because the vast majority of these events will simply fail to fix in the population, and thus be unobserved. Instead, we only wish to simulate events which survive to the present day and are observed in at least one sampled individual. (We use the terms ‘surviving’ and ‘observed’ interchangeably, as a lineage which survives in an unsampled individual is undetectable.) In order to do this, we must consider the probability of survival of each event.

In the MLMSC model, the survival probability is not constant. This can be seen by considering the following simple example: a duplication occurs ‘just before’ a speciation into two species leaves. In this case, the duplication can survive because it is observed in either of the descendant species (Figure 9a,b) and possibly in both (Figure 9c). Since the probabilities of being observed in either species are close to independent, the total survival probability is roughly twice that of a duplication which occurs in a terminal branch of the species tree (Figure 9d).

**Figure 9:**
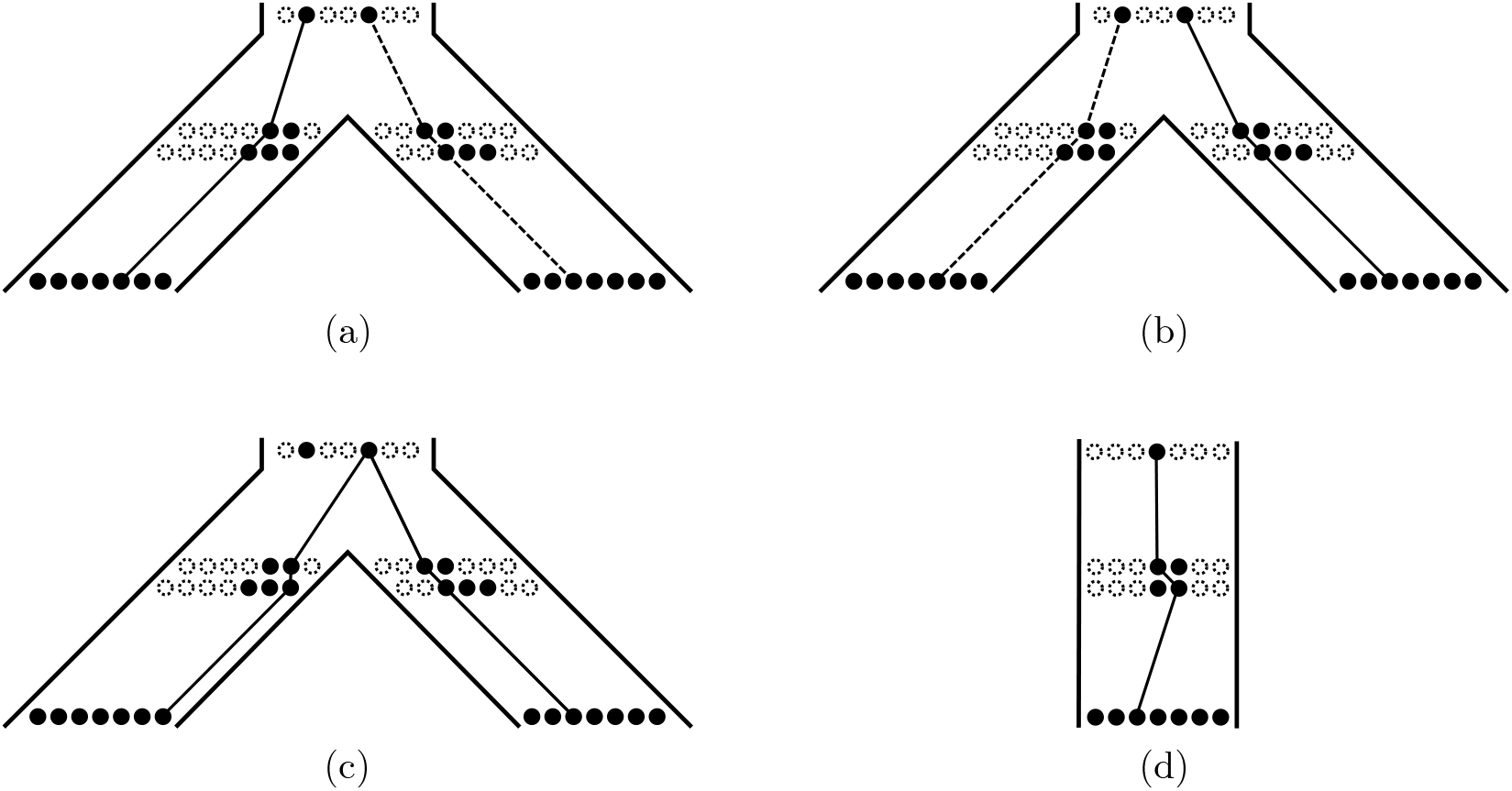
Although duplications occur at a constant rate, they do not survive with constant probability. (a-b) Duplication ‘just before’ a speciation. A duplication in either of the filled individuals will survive (we use the convention that the dashed line is not chosen as the haplotype tree). (c) Only a duplication in one of the individuals will survive, but this scenario requires a near-immediate coalescence and is unlikely. (d) Duplication in a species leaf. Only a duplication in one of the individuals can survive.

To simulate surviving duplication and transfer events at the correct rate, we introduce the *coalescent-rate process*. The simplest version of this process — applied to unlinked duplications — runs a coalescent process in the unilocus tree, and then simulates events at constant rate on the branches of the trees obtained in this way. The events are then considered to occur in the corresponding branches of the unilocus tree (see Figure 10). This allows us to simulate surviving duplications at the correct rate. Some modifications must be made for transfers and linked duplications, and we give further details in the Supplementary Material.

**Figure 10:**
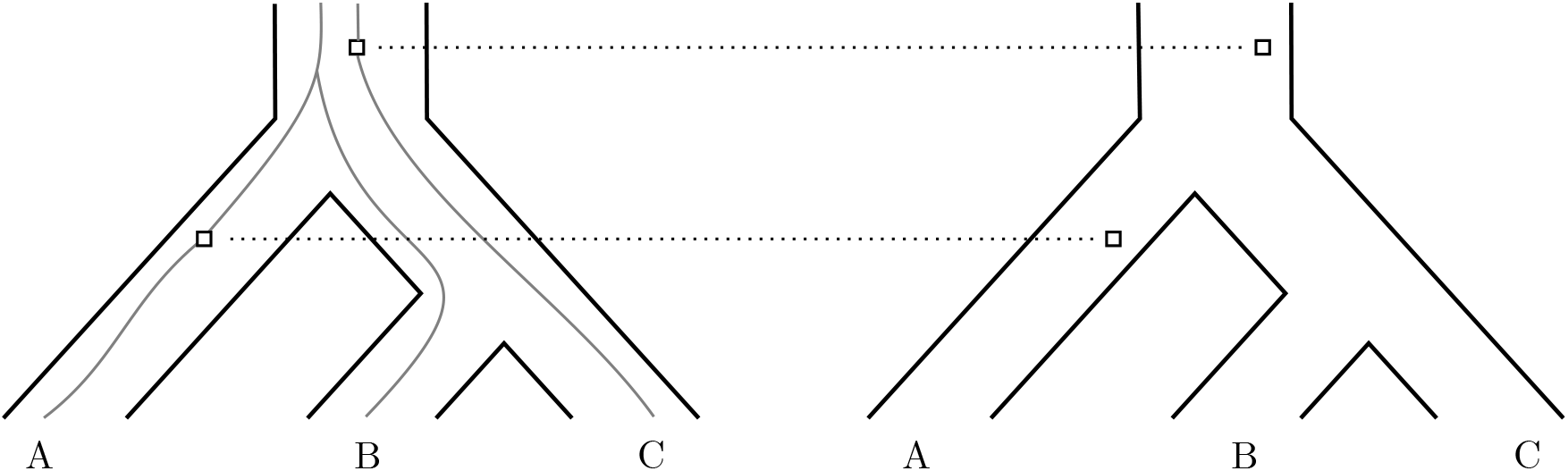
An example of the coalescent-rate process. Firstly, ‘temporary’ trees are sampled from the multispecies coalescent, and then events are sampled at constant rate on the branches of these trees. Finally, the ‘temporary’ trees are removed and the sampled events are considered to occur in the corresponding branches of the unilocus tree.

Unlike duplications and transfers, losses which are observed must occur on a surviving gene lineage, and thus they are sampled from the haplotype tree, instead of the unilocus tree, at a constant rate. Observe that this allows us to lose an allele which is not completely fixed in the population, resulting in CNH.

### Generating New Loci and Unilocus Trees

Once DTL events have been generated, the effect of each event is applied in a forwards-in-time order. The haplotype tree is truncated at each loss event. At each duplication or transfer event, a new locus is created, with a corresponding unilocus tree. The unilocus tree shows the evolution of all species which could possibly contain the locus (i.e., all descendants of the species where the locus is created). This is the subtree of the species tree *S*, starting from the time (and branch) of the creation of the locus (by duplication or transfer).

### Generating a Haplotype Forest — Unlinked Loci

Once we have generated the unilocus tree for a new locus, we then simulate the haplotype tree and forest. If the new locus is unlinked to the parent locus (i.e., it is created by transfer or unlinked duplication), then it evolves completely independently from its parent. To generate the haplotype tree and forest for such a locus, we introduce a new process called the *incomplete multispecies coalescent*; here, we do not require that all extant genes have to coalesce to their most recent common ancestor by the time of origination, i.e., at the root of the unilocus tree. Instead, we simply stop the coalescent process at the time of origination of the locus, thus producing a haplotype forest. One of these trees is then randomly chosen to be the haplotype tree. In this way, a duplicated/transferred gene copy does not have to be transmitted to all descendant species, resulting in CNH. An example of this process is shown in Figure 11.

**Figure 11:**
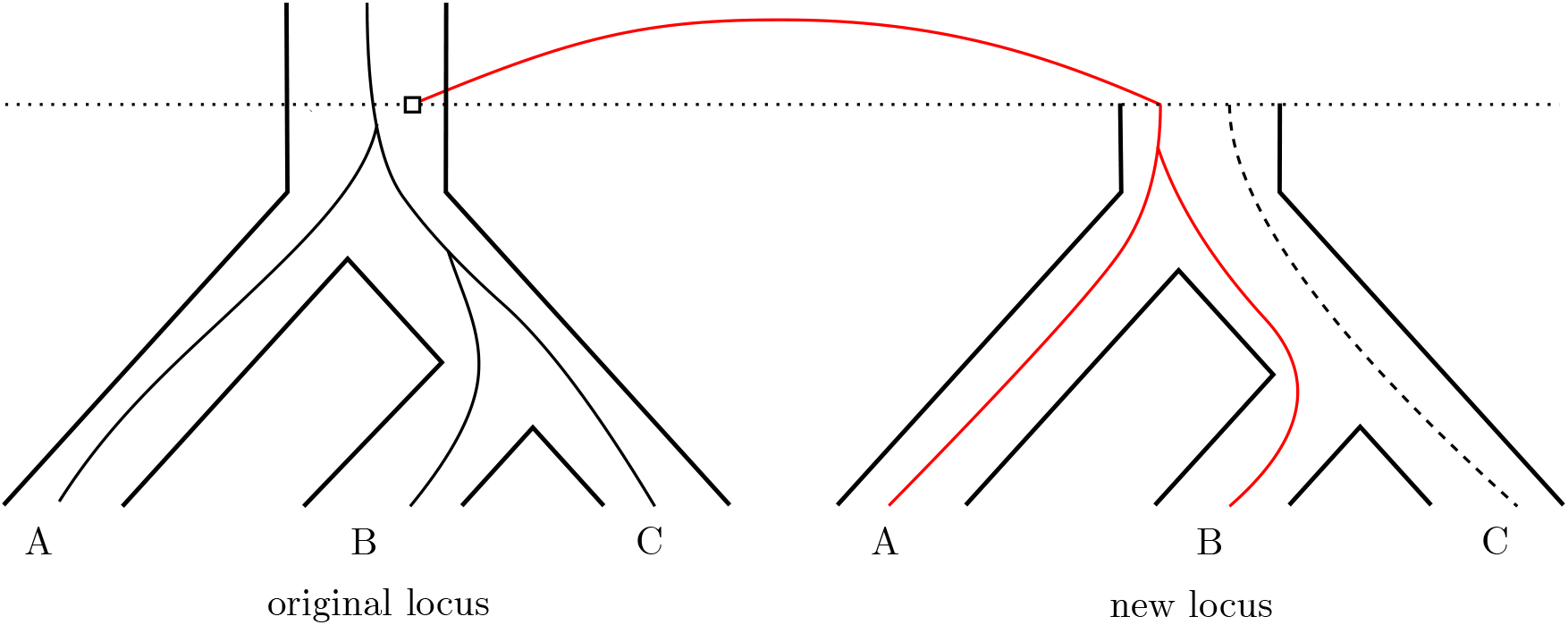
An example of the incomplete multispecies coalescent. Two trees (red and dashed) are generated in the new locus using the multispecies coalescent. The red tree is randomly chosen to be the haplotype tree for this locus, while the dashed tree is part of the haplotype forest but is not the haplotype tree. The root of the red tree represents the individual carrying the new copy.

### Generating a Haplotype Forest — Linked Loci

To generate the haplotype tree and forest in a locus created by linked duplication, we introduce a new process called the *linked coalescent*. In this process, the haplotype forest in the parent locus is mirrored into the new unilocus tree and used as a pre-existing genealogy which can be followed by the lineages in the new locus. The incomplete coalescent is then run, starting from a single lineage in each extant species which is coalesced with the pre-existing genealogy. Gene lineages which are coalesced with the pre-existing genealogy must follow it. In this way, the genealogy of the new locus depends on the genealogy of the existing locus; in the absence of recombination, the two will be identical.

To model the effects of recombination, lineages in the new locus which are coalesced with the pre-existing genealogy can ‘uncoalesce’ from it at a fixed rate, representing a recombination event between the new locus and its parent. In contrast, lineages in the new locus which coalesce with each other cannot uncoalesce, because they represent actual lineages. Lineages which are not coalesced with the pre-existing genealogy can coalesce with each other, or with the pre-existing genealogy, at rates consistent with the ordinary coalescent. An example of this process is given in Figure 12.

**Figure 12:**
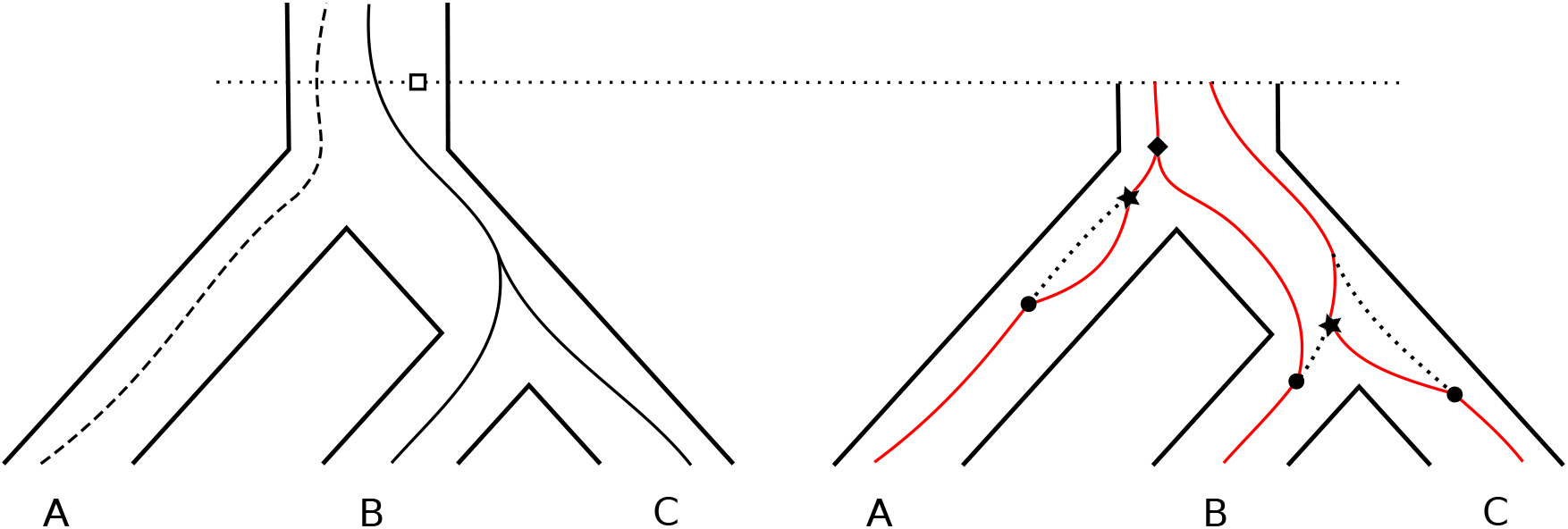
The linked coalescent. On the left, we have the original unilocus tree and its haplotype forest (the haplotype tree is solid and other trees in the haplotype forest are dashed); on the right, the new unilocus tree with a linked coalescent process run on it. The coalescent process starts linked with the genealogy copied from the parent locus in all species, then ‘uncoalesces’ due to recombination (black circles) in all species at different times, coalesces back to the mirrored genealogy (black stars) in *A* and the ancestor of *B* and *C*, and, finally, the lineages that descend to *A* and *B* coalesce with each other (black diamond).

This process produces the haplotype forest for the new locus, which may contain more than one tree. We choose one uniformly at random to become the new haplotype tree for that locus. This haplotype tree must then be (potentially) joined back to the haplotype tree in the parent locus, as detailed below.

### Assembling the Full Gene Tree

Once all events, unilocus trees, and haplotype trees have been generated, we assemble the full gene tree by concatenating each haplotype tree in a locus to the haplotype tree in its parent locus, starting from loci which have no descendants.

Of particular note is that a duplication may either be *ancestral* (the duplicating individual is a direct ancestor of a sampled individual in the genealogy of the parent locus), or *non-ancestral* (the lineage of the duplicating individual does not survive, or is not sampled, in the parent locus); see Figure 13. For unlinked loci, it can be assumed that only non-ancestral duplications occur, as the probability of survival in the parent locus, which is independent of the existence of the duplicated locus, is extremely small and can be safely ignored. However, when modelling linked loci, it is important to distinguish between these two cases.

**Figure 13:**
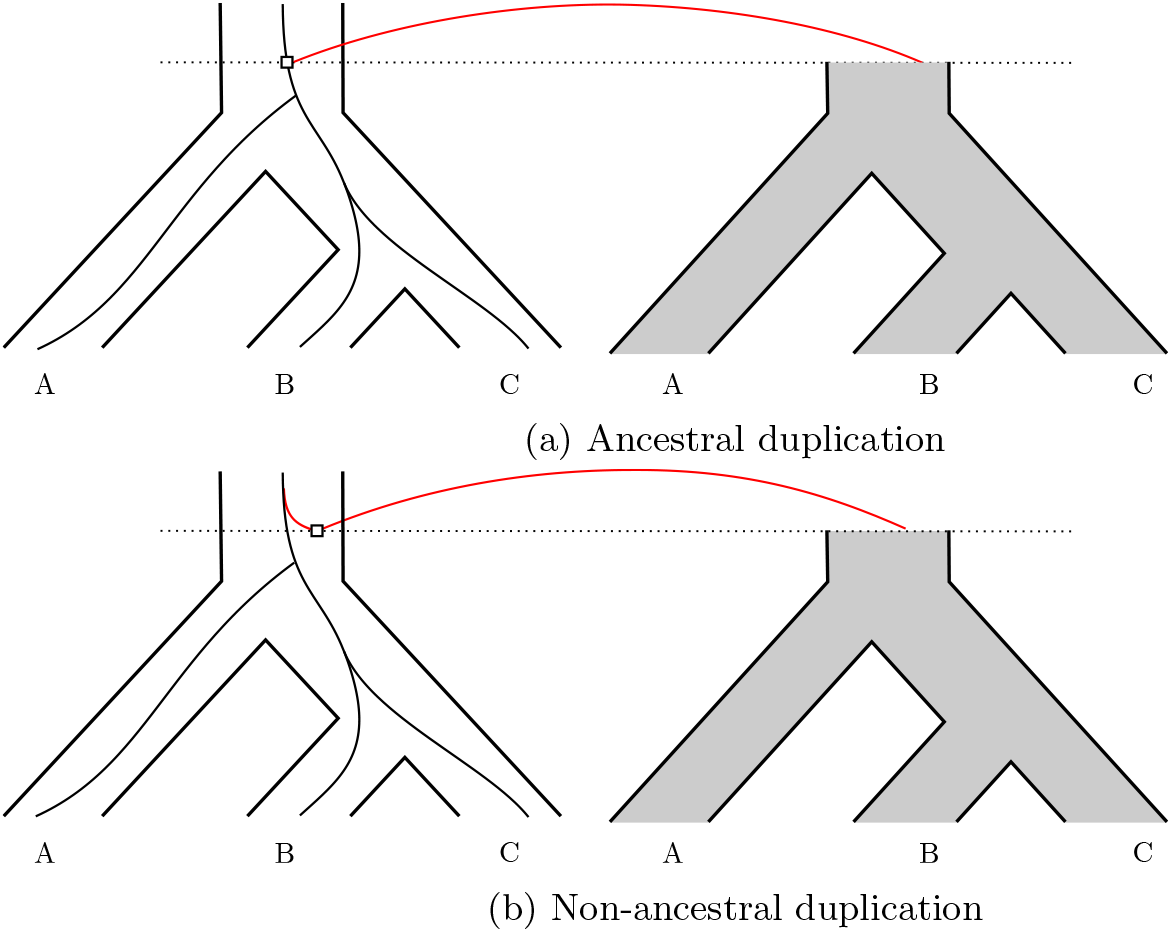
Ancestral vs non-ancestral duplications. (a) A duplication is ancestral if the duplicating individual is a direct ancestor of a sampled individual in the genealogy of the parent locus. (b) A duplication is non-ancestral if the lineage of the duplicating individual does not survive, or is not sampled, in the parent locus.

By associating lineages in the new locus with lineages in the parent locus, the linked coalescent provides a natural way to determine if a linked duplication is ancestral or non-ancestral. If the haplotype tree of a linked locus is coalesced (at the time of its creation) with a lineage copied from the parent locus, the duplication is ancestral. In this case, the haplotype tree of the child locus is joined directly to the lineage with which it is coalesced in the parent locus at the time of the event (Figure 13a).

On the other hand, if the locus is unlinked or if the haplotype tree is not coalesced with a copied lineage, the duplication is non-ancestral. In this case, we can treat the duplicating (or transferring) individual as another member of the population in the parent locus at the time of the event. We then follow this lineage backwards in time via the incomplete coalescent until it coalesces with the haplotype forest in the parent locus, or we reach the time of origination of the parent locus itself (Figure 13b).

Observe that this means that a locus can be entirely lost via sorting effects only. Consider the example in Figure 14: in a locus 1, originating at time *t*_1_, a duplication occurs at time *t*_2_, resulting in locus 2. Under the incomplete coalescent, it might be that the duplicating lineage fails to coalesce with any lineages in locus 1 by time *t*_1_. This can be interpreted as a duplication arising in an individual who does not actually contain any gene copy at that locus — i.e., an impossibility. In this case, the duplication is discarded and no new locus is created.

**Figure 14:**
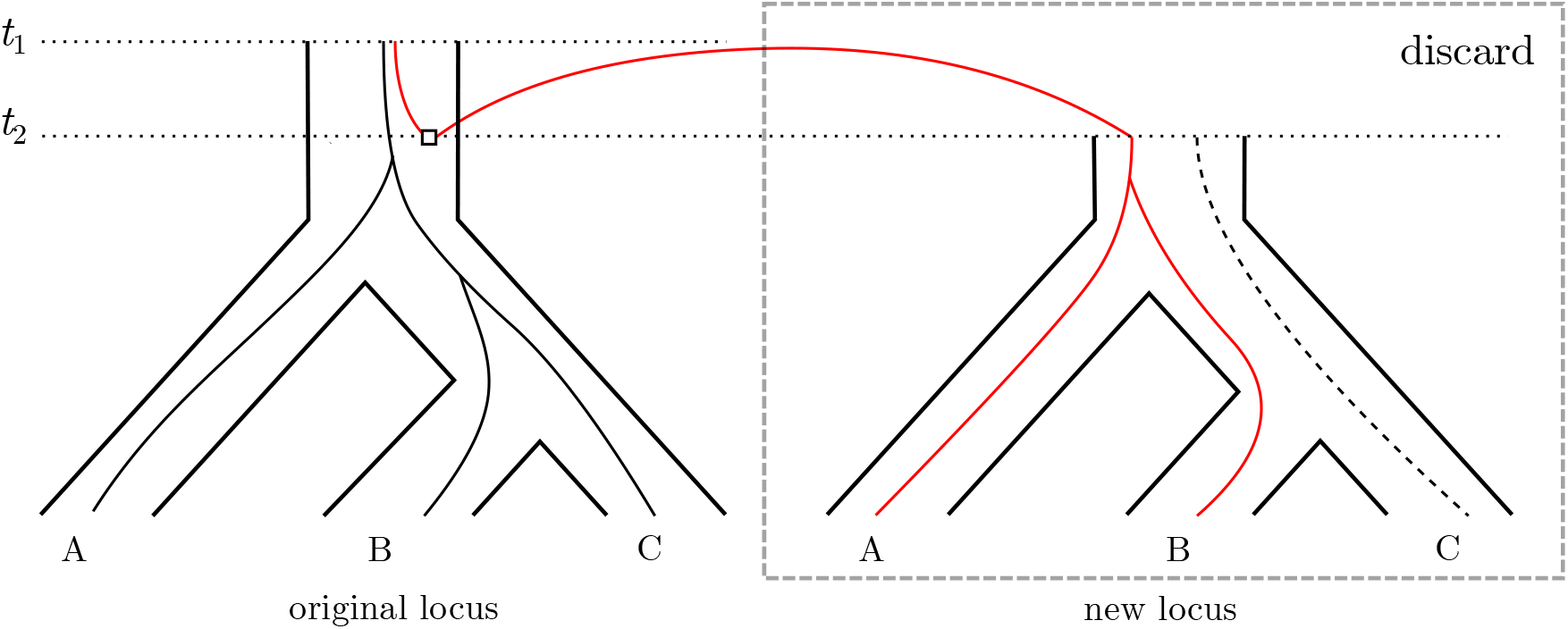
The new haplotype tree does not coalesce with the haplotype forest in the parent locus by the time of its creation, so the duplication is discarded. (Here we use the convention “dashed implies not chosen as haplotype tree”.)

When all haplotype trees have either been attached to the haplotype trees of their parents or discarded, we are left with one remaining tree starting in the original locus, which we take as the full simulated gene tree.

More details on the generating process, including a full example, formal notation, and pseudocode, are given in the Supplementary Material. We have implemented a simulator for this process, available at http:github.com/QiuyiLi/MLMSC.

## Model Comparison

In this section, we discuss the limitations of the current models in the literature, and show how the MLMSC model is subject to none of these limitations. Our discussion is summarised by, but not limited to, Table 1.

**Table 1:** Comparison of the models in the literature.

|  | Transfers | CNH | Recombination | Generating process | Reconciliation method |
| --- | --- | --- | --- | --- | --- |
| MLMSC | ✓ | ✓ | All | ✓ | × |
| DLCoal | × | × | Infinite | ✓ | ✓ |
| Haplotype tree model | × | × | None | ✓ | × |
| IDTL | ✓ | × | Limited | × | ✓ |
| Notung | ✓ | × | Infinite | × | ✓ |

### MLMSC vs DLCoal

The DLCoal model simulates losses on the locus tree, instead of the haplotype tree. Assigning a loss to a locus, instead of a branch of the haplotype tree, means that the loss of an allele due to lineage sorting cannot be modelled. Consider Figure 15a: here, two alleles were present in the ancestral population of *B*, *C*, and *D*, one of which was subsequently lost, i.e., replaced by the null allele. The null allele is then sorted into species *B* and *C*. The DLCoal model can only produce this gene tree by invoking at least two losses, as in Figure 15b. In order to properly capture this scenario, losses must be placed on the haplotype tree rather than the locus (or unilocus) tree, which is exactly what is done in the MLMSC model.

**Figure 15:**
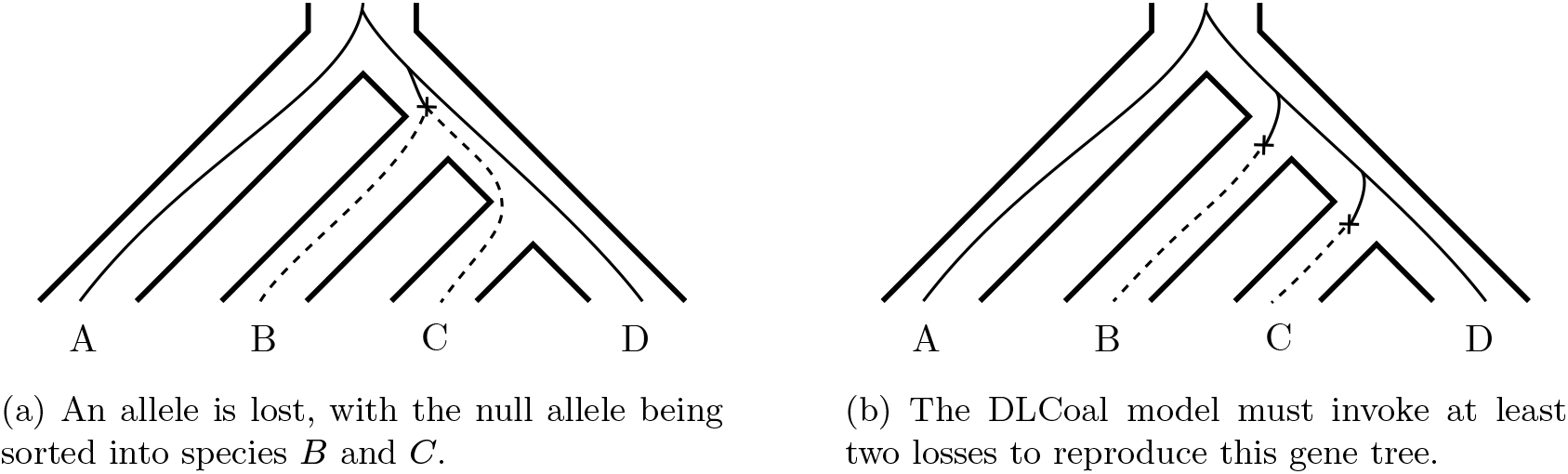
DLCoal cannot model lost alleles.

In a similar vein, the DLCoal model also assigns duplications to the locus tree. These duplications must then rejoin the haplotype tree via the multispecies coalescent; in other words, they are assumed to be non-ancestral. As discussed above, this is a reasonable consequence of the model assumption that all loci evolve independently (i.e., are unlinked), since the probability of an ancestral unlinked duplication is vanishingly small. However, in the more realistic case that some duplicated loci may be linked to their parent loci, the survival of a duplicated gene is correlated to the survival of its parent gene, and the probability of an ancestral linked duplication, knowing that the new duplicate exists, is non-negligible. Thus it is important to model both ancestral and non-ancestral duplications, as the MLMSC model does.

Finally, in the DLCoal model, a duplicated gene is either lost, or fixed in all possible descendant species. This means there can be no copy number hemiplasy. For example, DLCoal cannot model the scenario in Figure 16a: an additional loss is needed, as in Figure 16b. By use of the incomplete coalescent, the MLMSC model can model both scenarios.

**Figure 16:**
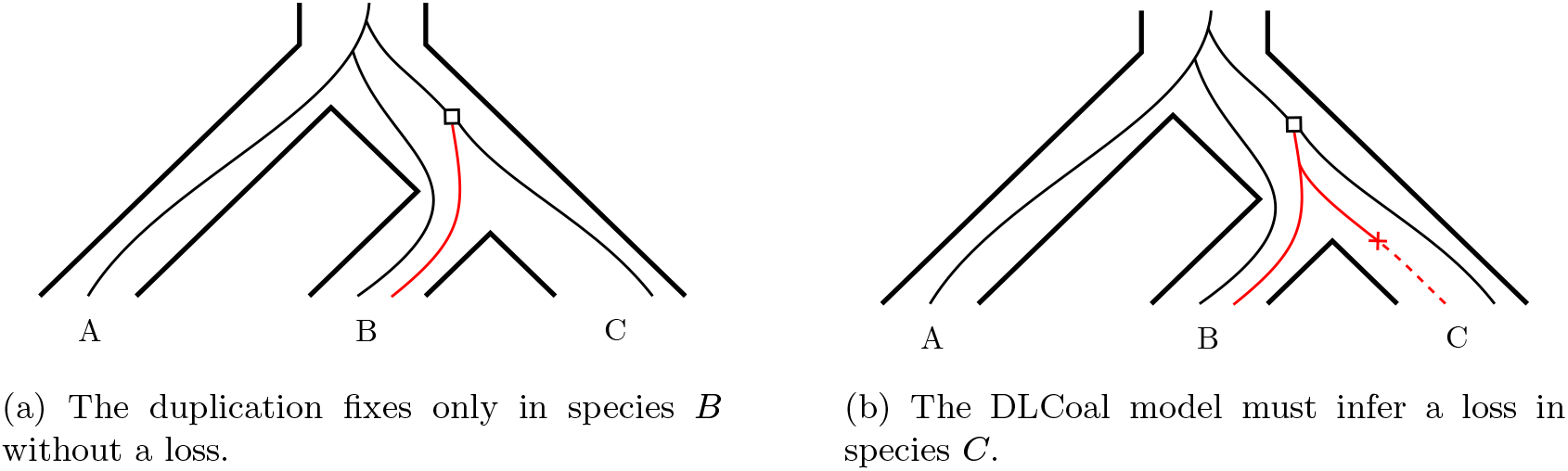
DLCoal cannot model copy number hemiplasy.

### MLMSC vs Haplotype Tree Model

In the haplotype tree model, a duplicated gene is assumed to have exactly the same genealogy as its parent gene; the model cannot model duplicated genes with different coalescent histories. For this to occur, we must assume that there is no recombination between loci, which is too restrictive. It is more realistic and flexible to allow the loci to be linked, where evolution in the loci are dependent but not necessarily identical, or unlinked, where evolution is completely independent. This is done in the MLMSC model.

Because the haplotype tree model applies the coalescent first, duplications and losses are assigned to gene lineages, rather than loci as in the DLCoal model. It is reasonable to apply duplications to gene lineages when there is no recombination (and thus only ancestral duplications can be observed). But, as discussed above, a more realistic model is to allow recombination, and model both ancestral and non-ancestral duplications, as done by the MLMSC model. On the other hand, the disadvantages of the DLCoal model do not apply to the haplotype tree model; for example, it can model both scenarios in Figure 15.

The haplotype tree model also does not fully allow for copy number hemiplasy; instead, keeping with the assumption of fully dependent loci, it enforces a restricted version wherein a duplicate must undergo the same coalescent process, and therefore be sorted into the same species as the parent gene. For example, the haplotype tree model cannot model the scenario in Figure 17a, where the duplicated gene is sorted into different species than the parent gene. It also cannot model the scenario in Figure 17b, where the duplicated gene undergoes a different genealogy from the parent gene. The MLMSC model can accommodate both of these scenarios, assuming either unlinked loci or linked loci with recombination.

**Figure 17:**
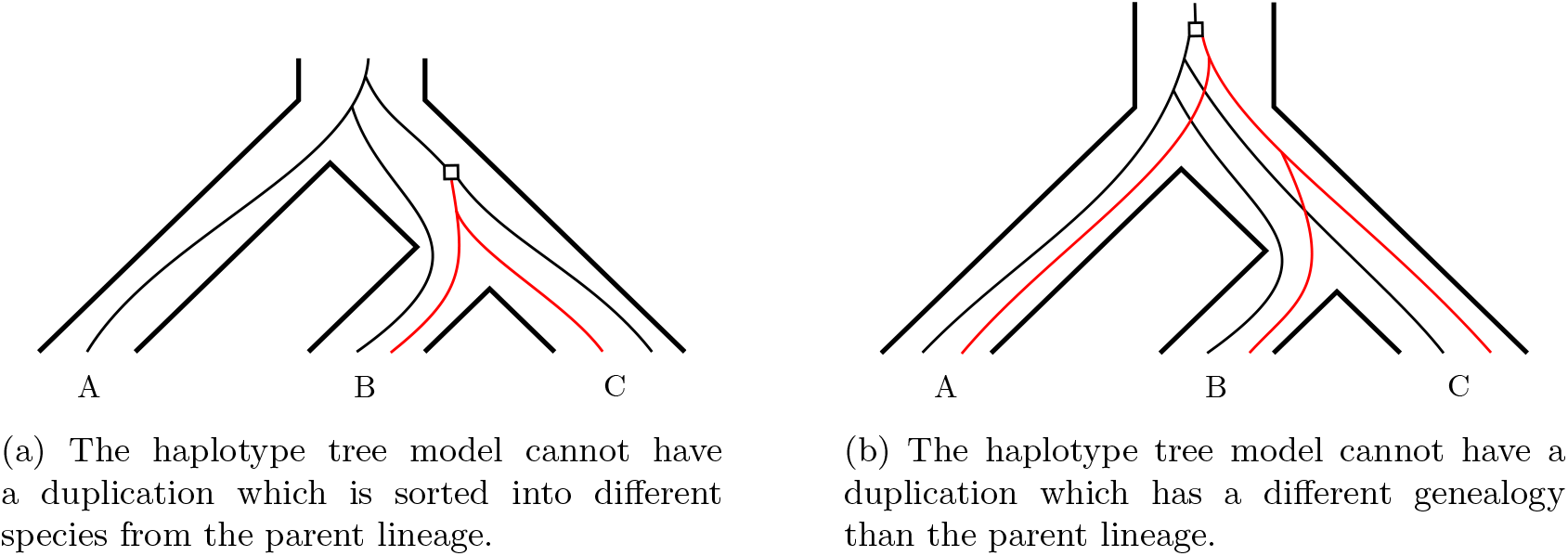
Limits of the haplotype tree model. Duplicated lineages are in red, while the parent lineages are in black.

### MLMSC vs IDTL Model

The IDTL model offers a kind of ‘halfway house’ between the DLCoal and haplotype tree models, where duplications, transfers, and losses are applied to gene lineages rather than loci, and duplicates can be sorted in different ways from their parent genes. However, some of the assumptions made in the model are computationally convenient but biologically questionable — for example, a duplicated gene must be sorted into the same species as its parent gene (i.e., no recombination allowed), but it may be sorted in a different way (i.e., recombination is allowed). For example, Figure 16a is also not allowed by the IDTL model. The MLMSC model is based explicitly on a model of the coalescent with recombination, and therefore incorporates recombination events in each species explicitly, allowing a greater range of scenarios.

### MLMSC vs Notung

As discussed above, Notung only allows ILS at a polytomy, and each possible sorting of genes at the polytomy is equivalent and equally likely; there is no ‘correct’ sorting which agrees with a specified species tree. In particular, this means that ILS is not penalised, and so any gene tree-species tree discrepancy which can be attributed to ILS is attributed to it, with the remaining discrepancies then explained by DTL. In contrast, MLMSC allows ILS everywhere in the tree, with probabilities based on the branch lengths of the species tree, and balances that with the DTL processes. It also specifies an underlying ‘true’ binary species tree which is always the most likely outcome for the gene sorting. Internal branch lengths in the MLMSC can be made arbitrarily short, which effectively covers what Notung represents by polytomies.

For computational convenience, the Notung model assumes that whenever a transfer occurs, the parent lineage must survive to the present day. This means that transfers must be ancestral. In reconciliation terminology, this means that there are no ‘transfer-loss’ events. As discussed above, this is not a realistic assumption, as only the transferred lineage needs to be observed; in fact, assuming selective neutrality, the chance of the parent lineage also surviving in the parent locus is 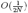, where 2*N* is the effective population size. The MLMSC model has no such restriction and can model transfer events accurately.

## Discussion

The MLMSC model generalises the multispecies coalescent to include duplications, transfers, and losses. By using the incomplete coalescent, haplotype forests, and disjoint unilocus trees, the MLMSC accounts for scenarios in which ancestral gene copy number polymorphisms are incompletely sorted among the descendant species, in contrast with existing models. By allowing for both linked and unlinked loci, the MLMSC model can also model more complex evolutionary scenarios than the existing models, while taking recombination into account in a natural way. Finally, the MLMSC recognises the fact, so far ignored, that the realised rate of duplication, transfer and loss becomes non-homogeneous if ILS is at work.

MLMSC is a particularly versatile model, which captures many evolutionary processes: speciation, gene duplication, gene loss, gene transfer, and genetic linkage. In this respect, MLMSC is more powerful than any of the existing models of gene family evolution (see Table 1). Note that what one can and cannot model has a strong impact on the accuracy of the estimation of species and gene trees, as pointed out for instance by (Boussau and Scornavacca (2020). We suggest that, in the presence of CNH, existing reconciliation methods might lead to biased estimates of gene trees, species trees, and/or evolutionary parameters, since every instance of a non-fixed duplication would incorrectly cost one loss if CNH is not explicitly accounted for.

Not all these processes, however, are necessarily relevant to every biological system. The prevalence of ILS, and consequently of CNH, depends on the ratio of effective population size to branch lengths (Pamilo and Nei, 1988), both of which vary by orders of magnitude among taxa and datasets. Scornavacca and Galtier (2017), for instance, suggested that ILS is only a minor determinant of conflicts between gene trees in phylogenetic analyses of the Mammalia clade, in which effective population sizes are relatively small (Romiguier et al., 2014) and branches typically represent millions of generations. DTL alone might be a sufficient model of gene family evolution at this scale. However, even in mammals, studies of closely related species have demonstrated the potential importance of ILS when short time frames are considered (Hobolth et al., 2011; Kutschera et al., 2014), suggesting that, as larger and larger data sets are generated and analysed, the pertinence of MLMSC should increase. Similarly, the modelling of genetic linkage between duplicates might be superfluous when relatively small and ancient gene families are considered, but should be crucial for the analyses of large and dynamic gene families, such as venom toxins (Fry et al., 2009) or olfactive receptors (Niimura et al., 2014), in which paralogous gene copies are often organized in clusters across the genome (Olender et al., 2020).

To assess the impact of CNH on gene evolution, we simulated gene trees on the fungal species tree used by Rasmussen and Kellis (2012), using three different duplication and loss rates (10^*−*10^, 5 *×* 10^*−*10^, and 10 *×* 10^*−*10^, duplication and loss rates are assumed to be equal), three effective population sizes (10^7^, 5 *×* 10^7^, and 10 *×* 10^7^), and 0.9 years per generation. For comparison, Rasmussen and Kellis (2012) used a duplication rate of 7.32 *×* 10^*−*10^, a loss rate of 8.59 *×* 10^*−*10^, an effective population size of 10^7^, and 0.9 years per generation. For each set of parameters, we ran 500 simulations and calculated the proportion of unilocus trees for which the haplotype forest contained more than one haplotype tree (i.e., CNH occurred). From Figure 18, we see that, for this data set, CNH is far from negligible, and as expected, its impact is higher when the effective population size is large — ranging from a bit less than 15% for *N_e_* = 10^7^, to more than 50% for *N_e_* = 10^8^. On the other hand, the proportion of CNH stays constant when varying the duplication and loss rate. This indicates that CNH is an important phenomenon which must be taken into account when modelling biological systems.

**Figure 18:**
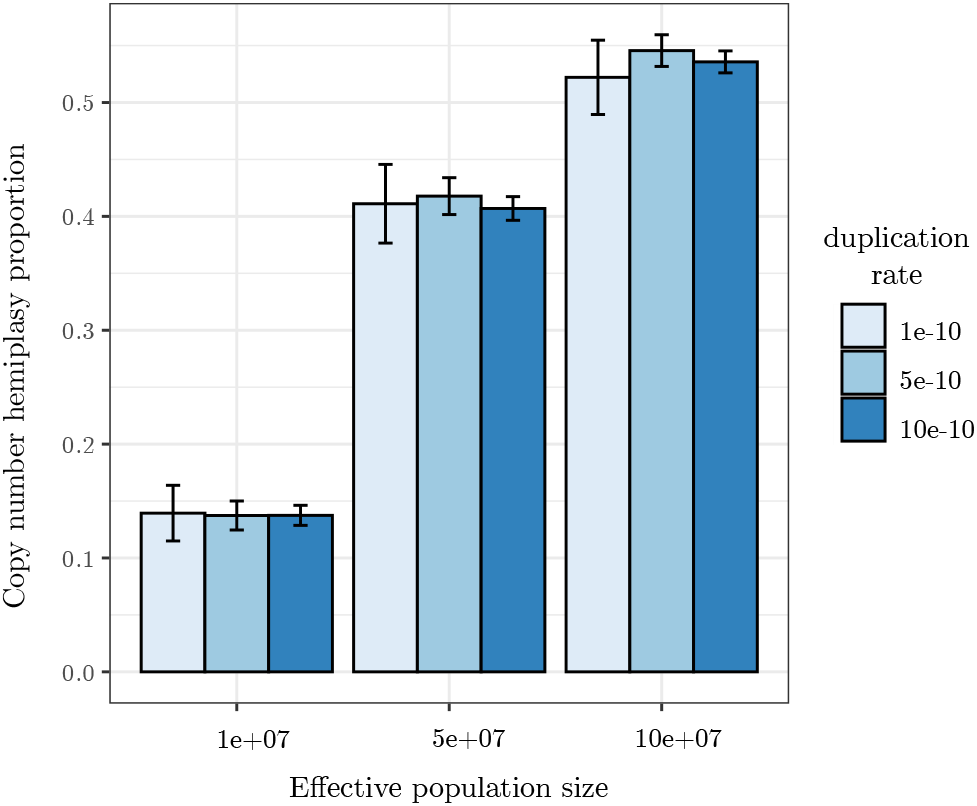
The proportion of CNH in our simulated data, varying the duplication and loss rate and the effective population size.

An implicit assumption in the MLMSC is that we can determine the linked or unlinked status of each locus solely with respect to its parent locus. In this respect, it is not fully cognisant of the linear structure of the chromosome. For example, if a locus (say locus 1) has two linked duplications in rapid succession (say loci 2 and 3), then one of the three loci must lie between the other two. If the order is (say) 1–2–3, then a recombination event between loci 1 and 2 must imply a recombination event between loci 1 and 3; this kind of dependency is not modelled by the MLMSC, which effectively represents the relations between linked loci as a tree rather than a linear structure. Moreover, in a scenario such as this, it may not be reasonable to assume that the recombination rate between loci 1 and 2 is equal to the recombination rate between 1 and 3. A full model which incorporates the linear structure of the chromosome would have to explicitly model the position of the gene copies, which we have elected not to do here for reasons of computational convenience.

We note that our goal here is not to correctly model recombination to its full extent, but to handle the case where the original locus and the duplicated one are next to each other and may thus have linked histories. In addition to the above, recombination is only allowed between (not within) duplicates. Traditional population genetic models, such as the coalescent with recombination and its approximations such as the sequentially Markov coalescent, permit more flexibility on this matter: recombination is allowed anywhere within a linear genome, and no pre-knowledge about the genealogy in neighbour loci is required. These models also have their drawbacks: all genomes are assumed to be aligned, with insertions and creations of new loci not allowed, and they assume that recombination occurs at a constant rate throughout the genome, ignoring the existence of hotspots or variation in rates due to gene function. One could say that our model does include this somewhat by considering a locus as a non-recombinant segment and only allowing recombination outside it.

Another limitation of our model is that it only accounts for a portion of the events shaping species and gene histories. For example, reticulated events such as hybridisation or reassortment, whole genome duplication events, and gene conversions are not modelled. Neither is “transfer from the dead” (Szöllősi et al., 2013), which is an important consideration when modelling transfers. There are no theoretical barriers to modelling this process, but it is unclear how to do so in a realistic and definitive manner. Two possibilities are to (a) maintain a separate ‘dead’ species which can only receive and donate transfers, or (b) allow transfers to ‘jump’ forward in time (i.e., allow the transfer origin to be selected at a time previous to the transfer target). For (a), we would need to set transfer rates to (and from) the ‘dead’ species; since we lack information about the structure of these species, a coalescent-rate process could not be used. For (b), we would need to determine the appropriate distribution of time between the transfer origin and target. Both of these models would go ‘outside’ the Wright-Fisher process which we are attempting to capture with the MLMSC model; thus, in order to maintain the elegance of our model, we have elected not to model this process here, but intend to include this in future work.

The analysis of gene families is an increasingly important component of comparative genomics (see e.g., Scornavacca et al., 2020, part 3). With so many fully sequenced genomes/transcriptomes available, the proportion of one-to-one orthologs in typical multi-gene, multi-species data sets will become smaller and smaller, and most of the information about the history of genomes and organisms must be extracted from multi-copy gene families. This is particularly true of organisms having large, complex genomes and undergoing frequent hybridisation and large-scale duplications, such as the economically important angiosperms (Glémin et al., 2019; Stull et al., 2020) and fish (Alda et al., 2019; Campbell et al., 2020), to name just a few.

In this context, the development of the general and arguably more realistic MLMSC model opens up promising research avenues. Simulations under the MLMSC can be used to assess the accuracy of current inference algorithms, and may help to confirm or contradict a number of published results. For instance, we suggest that neglecting ILS in phylogenetic analyses of gene families might bias the estimation of the timing of speciation/duplication. Patterns of gene loss subsequent to a whole-genome duplication, which is often interpreted in terms of evolution of gene function, could also be affected by ILS to an extent that remains to be quantified. Finally, in principle, the MLMSC model and its generating process could be used in an inference framework, i.e., serve as the basis for the development of new reconciliation algorithms, or parameter estimation methods (perhaps via Approximate Bayesian Computation). This would presumably be of great utility, while requiring substantial additional work.

## Supporting information

Supplementary material: full model specification

## Acknowledgements

YBC and CS acknowledge the Partenariat Hubert Curien (Hubert Curien Partnership) for providing grants for collaborative travel. CS was funded by grant ANR-19-CE45-0012 from the Agence Nationale de la Recherche. QL would like to acknowledge Geoffrey Law for providing assistance with programming.

