## Supplementary material: full model specification for "The Multilocus Multispecies Coalescent: A Flexible New Model of Gene Family Evolution"

### Full Model Specification

Here, we provide more formal details of the generating process of the MLMSC model. The process starts with the species tree as the unilocus tree for the original locus. Then we alternately generate a haplotype tree and forest for that locus, or new loci and unilocus trees, until all simulated duplication and transfer events are accounted for. The haplotype trees are then concatenated together to form the full gene tree.

For ease of reading, we provide a full example of these operations in Figures S1, S2, and S3. The formal statistical details are provided after the example. The pseudocode, to which we refer in the main text, is given in Algorithms 1 and 2 at the end of the section.

#### 1. Generating a Haplotype Tree for the Original Locus

We start with a species tree $S$ (line 1, Algorithm 1). In the original locus (which we denote as locus 0), the unilocus tree $S_0$ is the original species tree $S$. For this locus only, the haplotype tree $G_0$ is generated according to the standard multispecies coalescent, starting from a single copy of the gene in each leaf of the tree (line 2, Algorithm 1). We also set the haplotype forest $\mathcal{G}_0$ to be the set $\{G_0\}$ (line 3, Algorithm 1).

For simplicity, we assume (here and in all further coalescents) that the effective population size $2N$ is constant over time and across species. The process can easily be generalised to more variable cases, with the caveat that the population size is a property of the species and must therefore remain the same across loci.

For loci created by duplication or transfer, a more complex process is required (line 4, Algorithm 1). We first describe how to generate new loci and unilocus trees, then return to generating haplotype trees within those unilocus trees.

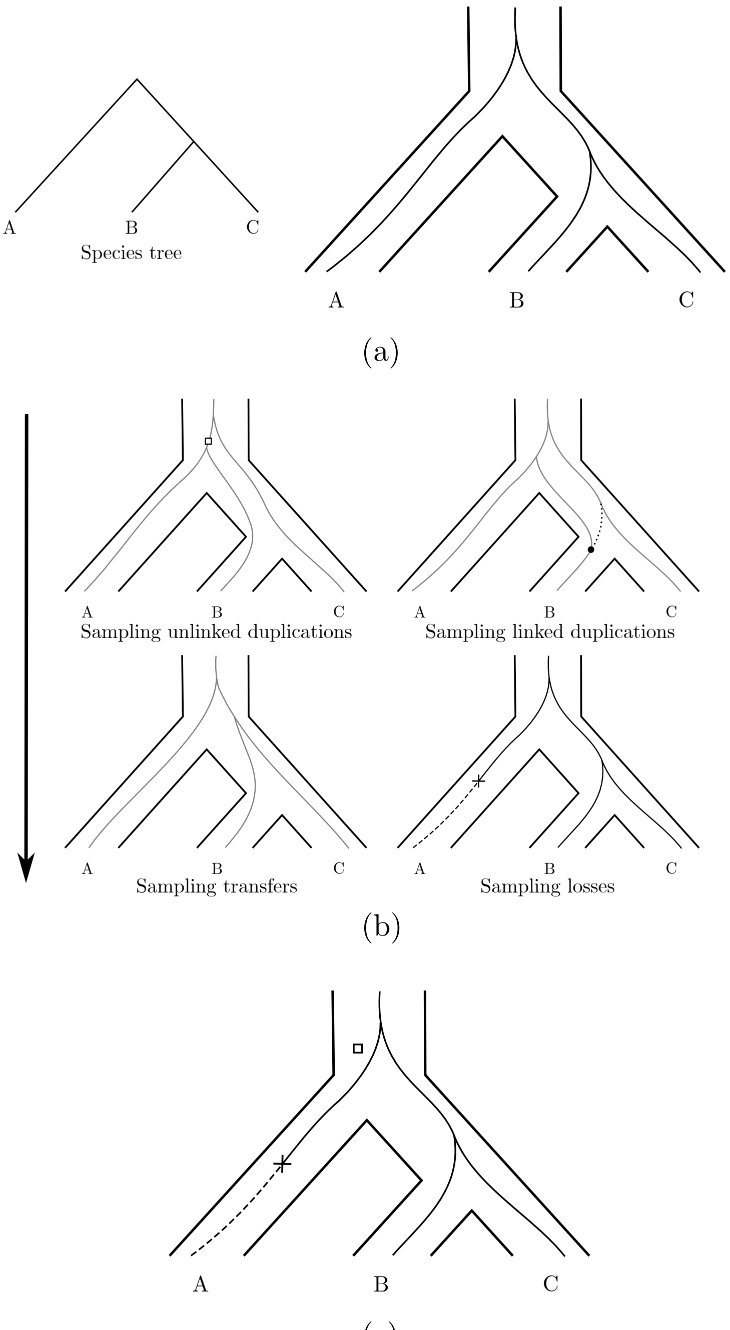


Figure S1: Modelling gene evolution in the original unilocus tree. (a) The species tree (top left) and original unilocus tree ('tubes', on top right), within which a haplotype tree (finer lines) is generated according to the standard multispecies coalescent. (b) Simulating unlinked duplications, linked duplications and transfers in the unilocus tree with coalescent-rate processes, and losses on the haplotype tree. Here, only one unlinked duplication and one loss are simulated, while no linked duplication or transfer is simulated. (c) The unilocus tree is decorated with the unlinked duplication, and the haplotype tree is truncated at the loss.

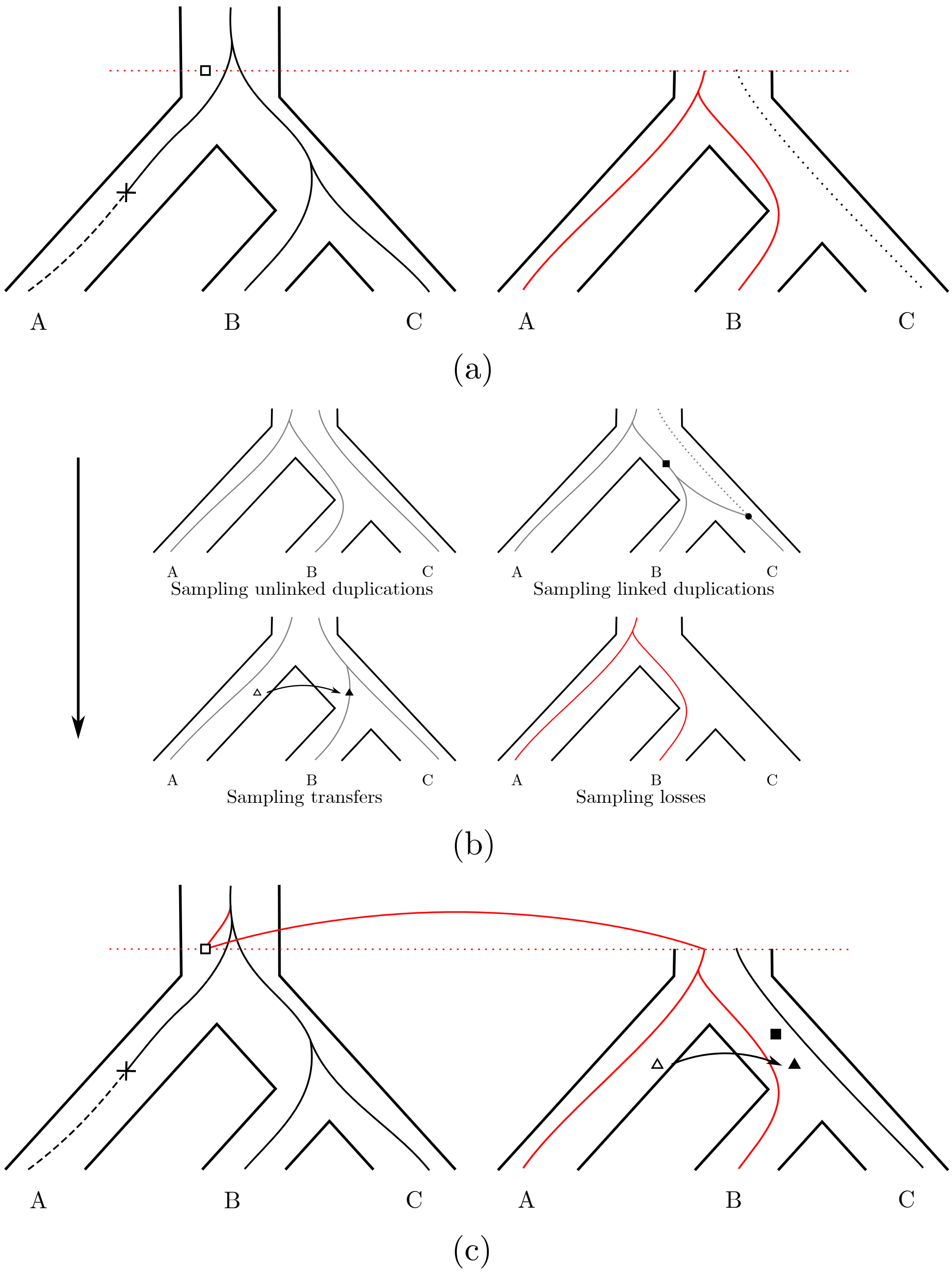


Figure S2: Modelling gene evolution in the first child locus. Here, we follow the duplicated gene in Figure S1. (a) We generate the new unilocus tree ('tubes', on the right) by copying the original unilocus tree ('tubes', on the left) from the time of the duplication. The new haplotype forest (finer lines, on the right) is generated according to the incomplete multispecies coalescent; the red tree is chosen to be the new haplotype tree. (b) Simulating events in the new unilocus tree, as in Figure S1b. Here, only one transfer and one linked duplication are sampled, while no unlinked duplication or loss is sampled. (c) The new haplotype tree is joined to the haplotype tree in the original locus, and the new unilocus tree is decorated with the simulated events.

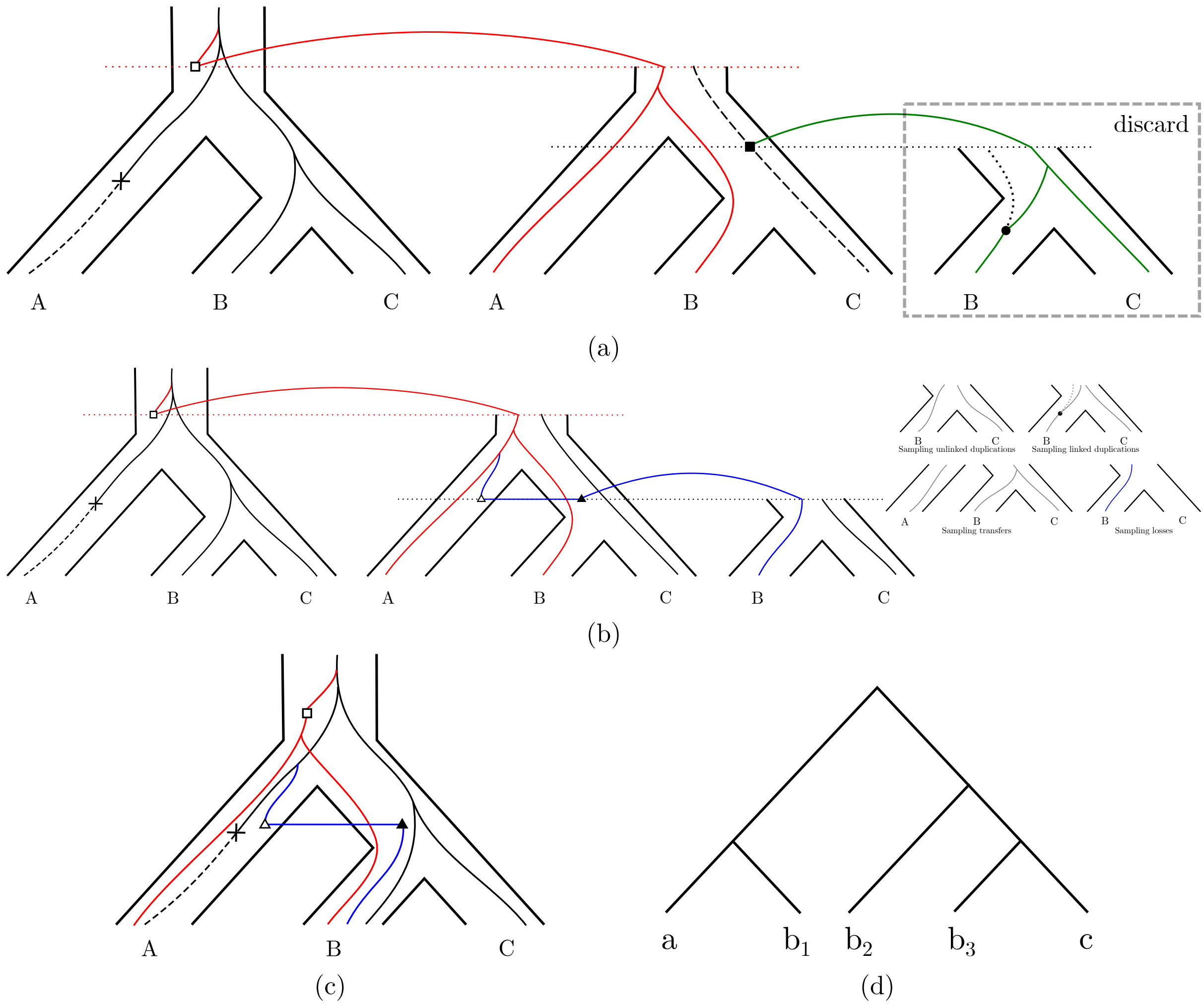


Figure S3: Modelling the evolution of the loci derived from the first child locus. Here, we follow the duplicated gene (a) and transferred gene (b) in Figure S2. (a) We generate the new unilocus tree ('tubes', on the right) by copying the parent unilocus tree ('tubes', on the centre) from the time of the duplication. The new haplotype tree (finer lines, on the right) is generated according to the linked multispecies coalescent. This tree is linked, in the parent locus, to a tree in the haplotype forest which is not the haplotype tree, and is thus discarded. (b) We apply a similar procedure for the transferred gene, which joins the parent haplotype tree and is thus kept. The new event sampling procedure (far right) does not simulate any new event. This concludes our example. (c) The final gene tree within the species tree. (d) The final gene tree.

### 2. Simulating Events

Suppose that we are simulating events from a locus $l$, with unilocus tree $S_l$, haplotype tree $G_l$, and haplotype forest $\mathcal{G}_l$. We are given a set of parameters $\boldsymbol{r} = (r_d, r_t, r_l, r_r, p_u)$, where $r_d$, $r_t$, and $r_l$ are the rates of duplications, transfers, and losses respectively, $r_r$ is the rate of recombination for linked duplications, and $p_u$ is the (fixed) probability that a duplication will be unlinked. The trees have branch lengths given in coalescent units, and by convention the present day is time $t = 0$, with $t$ increasing as we go backwards in time. The rates $r_d, r_t, r_l$, and $r_r$ are given in units of events per individual per locus per coalescent unit.

Recall that we only wish to model surviving DTL events, not all of them. A duplication may either be ancestral (the duplicating individual is a direct ancestor of a sampled lineage in the parent locus), or non-ancestral. Either of these kinds of duplication can survive, and so we can consider a duplicating individual to be any one of the population, and thus sample duplications from the unilocus tree. Likewise, (surviving) transfers can also be sampled from the unilocus tree.

On the other hand, a loss in a lineage which does not survive to the present day will not be observed; in other words, losses are always ancestral. Hence losses should be sampled from the haplotype tree at constant rate $r_l$ (line 4, Algorithm 2). The constant rate is justified because a population which contains multiple alleles will lose each of them at the same rate as a completely fixed gene, and thus the overall loss rate will be proportional to the number of existing lineages.

In order to sample surviving duplications and transfers, we must consider both the rate at which the events occur, and their probability of being observed. We assume that the events (all events, not surviving events) occur at a constant rate per individual per locus per coalescent unit, as given by the parameters. Thus the overall rate of each event across a population is $2N$ multiplied by the per-individual rate. We then need to scale this rate by the probability of survival of the event. This probability differs for unlinked duplications, transfers, and linked duplications. We consider each in turn.

#### 2.1. Simulating unlinked duplications (line 1, Algorithm 2)

Note that in the DLCoal model, because no CNH is allowed, the survival probability of a duplication is constant. Since unlinked duplications occur at a constant rate, surviving unlinked duplications can also be simulated at a constant rate. In the MLMSC model, we use the incomplete coalescent to model CNH, so this property no longer holds.

In the MLMSC model, unlinked duplications occur at a constant rate of $2Np_ur_d$. To calculate the survival probability of an unlinked duplication at a locus $l$, consider a duplication which occurs at a certain time and species in $S_l$, and consider the sampled individuals in the extant descendants of this species. Each of these individuals will have a single direct ancestor for locus $l$ at the time of the duplication, which may or may not be distinct from each other. The duplication will be observed if and only if one of these ancestors is the duplicating individual, in a population of size $2N$. The probability that such a duplication is observed is therefore $\frac{1}{2N}$ multiplied by the expected number of lineages at the time of the duplication, under the multispecies coalescent.

It is difficult to obtain a closed formula for this number, but there is a simple way to simulate at the correct rate. To do this, we run a (incomplete) multispecies coalescent within $S_l$, and then sample events with a constant-rate Poisson process at rate $p_ur_d$ from the branches of the resulting coalescent trees. These events will be considered as duplications at the corresponding branch and time in $S_l$. We call this a *coalescent-rate process*; an example of this process is given in Figure S4.

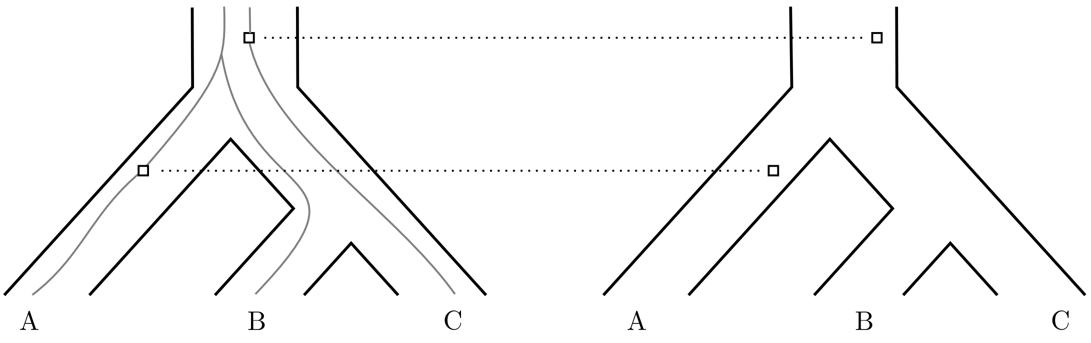


Figure S4: An example of the coalescent-rate process. Firstly, 'temporary' trees are sampled from the multispecies coalescent, and then events are sampled at constant rate on the branches of these trees. Finally, the 'temporary' trees are removed and the sampled events are considered to occur in the corresponding branches of the unilocus tree.

At any time, the coalescent trees will, on average, trivially have the expected number of lineages under the coalescent, and so the resulting rate is correct. We must also specify a rate for the coalescent-rate process, which is the rate at which events are sampled from the branches of the coalescent trees (in this case $p_u r_d$).

#### 2.2. Simulating transfers (line 2, Algorithm 2) —

It is important to note that the coalescent-rate process reflects the probability of a gene lineage surviving if it *appears* (by duplication or transfer) in a particular species. A duplication always appears in the same species that it originates from, but a transfer does not. Thus the coalescent-rate process must be used to select the target species of the transfer, not its origin species. Additionally, while a transfer from locus $l$ must originate from within $S_l$, it need not appear in $S_l$, but could appear in any contemporary species different from the origin species.

In order to simulate surviving transfers from locus $l$, we thus perform a coalescent-rate process with rate $r_t$ over the *entire* species tree $S$. Each resulting event marks the target species of a transfer, and we subsequently choose an origin species uniformly at random from *all* other contemporaneous species. If this origin species does not lie in $S_l$, the entire transfer is discarded. An example of this process is given in Figure S5.

In practice, it is less efficient to simulate many transfers and discard some, so we take advantage of the fact that transfers originate from each species at an equal rate, since we assume that all population sizes are equal. Hence, we scale the overall transfer rate to each species by the proportion of possible origin species which are also contained in $S_l$, and then select an origin species uniformly at random from the possible origin species in $S_l$. It is easy to see that this is equivalent to the process described above, but without having to discard some simulated transfers.

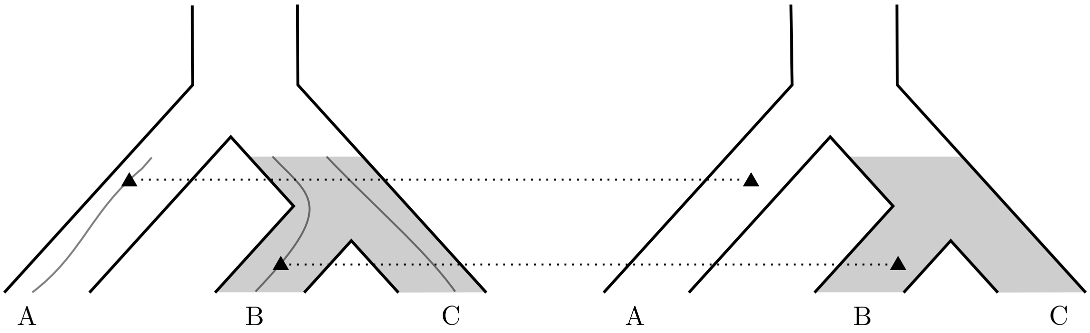


(a) Transfer targets (represented by black triangles) are generated using a coalescent-rate process over the entire species tree: an incomplete coalescent process is run on the species tree, and transfer targets are sampled on the coalescent trees generated in this way (left). The coalescent trees are then discarded and the transfer targets kept (right).

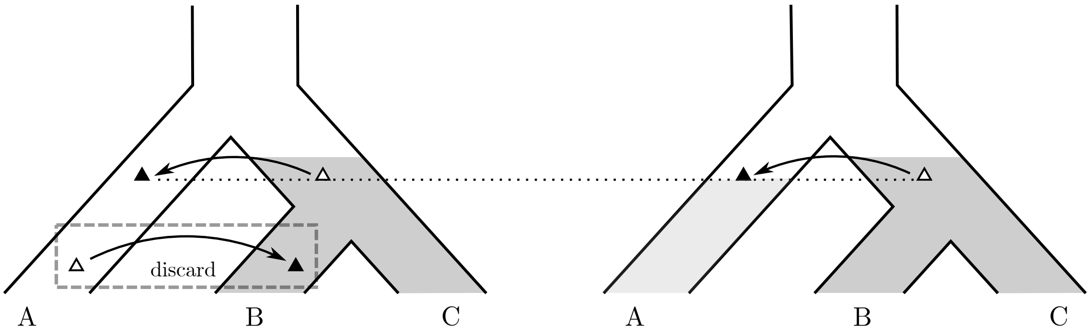


(b) For each transfer target, we simulate a transfer origin (represented by a white triangle, left). Transfers originating from outside the unilocus tree (depicted as shaded), e.g., the lower transfer in our example, are discarded (right).

Figure S5: Simulating transfers from a given unilocus tree (with leaves $B$ and $C$, shaded) by first simulating targets (a) and then origins (b). The transfer must originate from inside the unilocus tree, since we are simulating transfers *from* it. In (b), the new unilocus tree derived from the only retained transfer (with leaf $A$) is shown in a lighter shading on the right.

#### 2.3. Simulating linked duplications (line 3, Algorithm 2) —

In a linked locus created by duplication, the probability of the duplication surviving is dependent on the genealogy of the parent locus. However, the principle is the same: a duplication will survive if and only if it is the direct ancestor of a sampled lineage in the new locus. The linked coalescent (described in Section "*Generating a Haplotype Forest — Linked Loci*") directly models the genealogy in such a locus, and thus it can be substituted in place of the ordinary coalescent when generating linked duplications.

To simulate surviving linked duplications originating from locus $l$, we therefore run a linked coalescent in the unilocus tree $S_l$, allowing the lineages to coalesce and uncoalesce (through recombination) with the existing haplotype forest $\mathcal{G}_l$. We then sample linked duplications as events at constant rate $(1 - p_u)r_d$ on the branches of the resulting coalescent forest. These events are then interpreted as linked duplications at the corresponding branch and time inside $S_l$. We refer to this as a *coalescent-rate process in the presence of* $\mathcal{G}_l$.

#### 2.4. Duplication and transfer rates —

Before continuing, we comment briefly on duplication and transfer rates. For a duplication/transfer to 'survive', there are three things which need to happen: 1. the event occurs; 2. the duplicated/transferred individual has a copy of the gene at the locus; and 3. we observe a descendant of the duplicated/transferred individual.

We have a fixed rate for point 1 (i.e., the rates $r_d$ and $r_t$); the coalescent-rate process handles point 3 by incorporating the probability to observe a descendant of the duplicated/transferred individual; and point 2 is accounted for by rejection sampling, i.e., potentially discarding loci when joining the haplotype tree of the new locus with that of the original locus (we will discuss this later, see Figures S10b, S11b and S11c). This implies that the 'actual' rates (i.e., the rates of surviving events which duplicate or transfer a copy of the gene) will be lower than the rates $r_d, r_t$. The actual relation between these pairs of rates is highly non-trivial.

### 3. Generating New Loci and Unilocus Trees (lines 11–12, Algorithm 2)

When a new locus is created, we create a unilocus tree for it, which is the subtree of the species tree starting from the time (and branch) of the creation of the locus. We then generate duplication, loss, and transfer events. In order to formally describe this process, we first introduce some notation. Given a tree $T$, $V(T)$ and $E(T)$ are the sets of their nodes and edges (branches) respectively. For a node $v \in V(T)$, $e_v$ represents the unique edge in $E(T)$ which has $v$ as its target node (the node at the bottom of the edge).

Now, given a unilocus tree $S_l$ and the corresponding haplotype tree $G_l$ and haplotype forest $\mathcal{G}_l$, we simulate losses as described above, producing a sequence of events $\boldsymbol{e^l} = \{(b_i, t_i)\}$, where $b_i \in E(G_l)$, and $t_i \geq 0$ is the time of occurrence of the loss (how the haplotype tree and forest are generated is described in the next section). Note that we must have $t_2 < t_i < t_1$, where $t_1$ and $t_2$ are the times of the top and bottom nodes of branch $b_i$ respectively. At each event, we decorate $G_l$ at that branch and time. An example of this process is shown in Figure S6.

We then simulate (as described above):

1. unlinked duplications in $\boldsymbol{e^{ud}} = \{(b_i, t_i)\}$, where $b_i \in E(S_l)$;
2. transfer targets in $\boldsymbol{e^t} = \{(b_i, t_i)\}$, where $b_i \in E(S)$;
3. linked duplications in $\boldsymbol{e^{ld}} = \{(b_i, t_i)\}$, where $b_i \in E(S_l)$.

In all cases, the same constraints on $t_i$ given for losses apply. At each duplication, we decorate $S_l$ at that branch and time, and at each transfer, we choose and decorate the origin branch of $S_l$ at that time. An example of the generation of unlinked duplications is given in Figure S7.

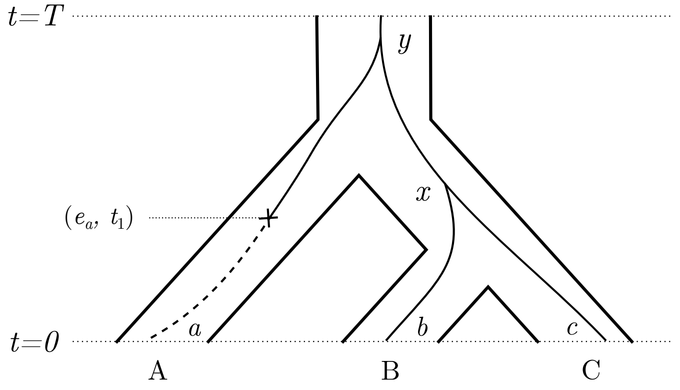


Figure S6: Losses are sampled from the haplotype tree with constant rate. $A$, $B$, and $C$ are species labels, while $a$, $b$, and $c$ are gene labels. Here, we have $\boldsymbol{e^l} = \{(e_a, t_1)\}$.

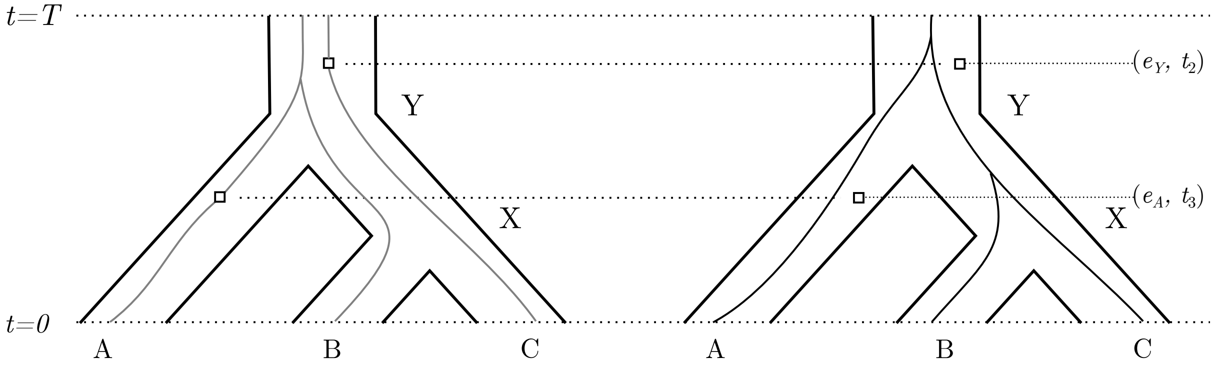


Figure S7: Duplications are sampled with a coalescent-rate process on the unilocus tree. Firstly, 'temporary' trees (in grey) are generated by running a multispecies coalescent within the unilocus tree. Then duplications are sampled at constant rate on the branches of the 'temporary' trees. Finally, the 'temporary' trees are removed and the sampled duplications are considered to occur in the corresponding branches of the unilocus tree (with the haplotype tree in black). Here, we have $\boldsymbol{e^{ud}} = \{(e_Y, t_2), (e_A, t_3)\}$.

Now, the effect of each event is applied in a forwards-in-time order:

- At each loss event, the haplotype tree is truncated on branch $b_i$ at time $t_i$, with further events on the same or descendant branches of $G_l$ (but not $S_l$) having no effect. (Note that the haplotype forest is *not* truncated.)
- At each duplication event $(b_i, t_i) \in \boldsymbol{e^{ud}} \cup \boldsymbol{e^{ld}}$, a new locus $m$ is created, with a unilocus tree $S_m$ which is a copy of the subtree of $S_l$ which starts at time $t_i$ on the species branch $b_i$ (see Figure S8).
- Likewise, at each transfer event $(b_i, t_i) \in \boldsymbol{e^t}$, $b_i$ represents the target of the transfer, not the source (as discussed above). The new locus will then have a unilocus tree created, which is a copy of the subtree of $S$ starting from branch $b_i$ at the transfer time $t_i$.

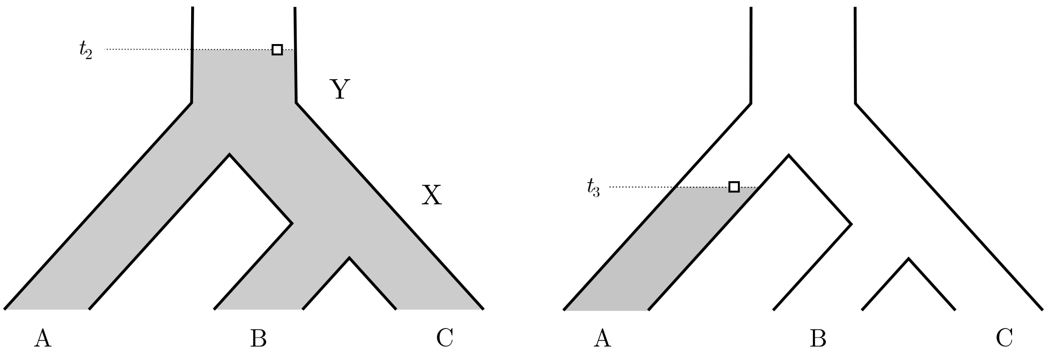


(a) For the duplication at time $t_2$, the new unilocus tree is the parent unilocus tree with the branch $e_Y$ slightly shorter, i.e., starting at time $t_2$ instead of $T$.

(b) For the duplication at time $t_3$, the new unilocus tree is the leaf branch $e_A$ of the parent unilocus tree, starting at time $t_3$.

Figure S8: The new unilocus trees generated by the events in Figure S7: (a) a duplication at $(e_Y, t_2)$ and (b) a duplication at $(e_A, t_3)$. The unilocus trees are depicted as shadings on the species tree, with height equal to the duplication time.

### 4. Generating a Haplotype Forest (lines 13–19, Algorithm 2)

Once we have generated the unilocus tree for a new locus, we must then simulate the haplotype tree and forest. If the new locus is unlinked to the parent locus (i.e., it is created by transfer or unlinked duplication), then the two loci evolve completely independently. We then run the multispecies coalescent within the unilocus tree, and stop it at the time of creation of the locus. We refer to this process as the *incomplete coalescent.* For a new locus $m$, this produces a collection of trees, which we take as the haplotype forest $\mathcal{G}_m$. A single tree from $\mathcal{G}_m$ is then selected uniformly at random as the haplotype tree $G_m$ (lines 14–15, Algorithm 2).

Suppose a locus $l$ has a linked duplication, creating a new locus $m$. We need to consider that the genealogies of the two loci are dependent, and take this into account when constructing the haplotype tree and forest (lines 17–19, Algorithm 2). We have some genealogical information from the parent locus $l$, represented by the haplotype forest $\mathcal{G}_l$.

Since $S_m$ is a copy of a subtree of $S_l$, we can identify all subtrees of the haplotype forest $\mathcal{G}_l$ which lie in the subtree of $S_l$ corresponding to $S_m$. Formally, if locus $m$ is created from an event $(b_i, t_i)$, then we take all subtrees of $\mathcal{G}_l$ which lie in the species branch $b_i$ and its descendants, truncated so that they start at time $t_i$. These subtrees are then copied to the new locus $m$ (see Figure S9b for an example).

We now run the *linked multispecies coalescent* in the new unilocus tree $S_m$. We start from a single lineage in each extant species, as with the ordinary multispecies coalescent. Because there is only one sampled individual per population, the lineage in each species begins coalesced to the copied subtree which has that species as a leaf. By construction, each extant species has one leaf in $\mathcal{G}_l$, so there is always exactly one possibility for this.

The coalescent then proceeds backwards in time, where the lineages which are coalesced with a copied subtree follow the same genealogy as that subtree. In particular, lineages which are coalesced with copied subtrees will coalesce with each other when the corresponding subtrees coalesce in the parent locus. However, with a constant rate $r_r$, each lineage which is coalesced with a copied subtree may have a recombination event (i.e., a recombination occurs between the two loci). If there are no recombination events, the haplotype forest in the new locus will be identical to that of the parent locus (Figure S9a).

When a recombination event occurs, the lineage in the new locus becomes 'uncoalesced' from the copied subtree, representing the fact that the individual in the population had different ancestors for the two loci. This lineage now becomes 'free' (not coalesced with a copied subtree) and is also followed backwards in time, where it may coalesce with a copied subtree of $\mathcal{G}_l$, or with another lineage in the new locus (which may itself be coalesced with a copied subtree). Note that lineages in the new locus $m$ which coalesce with each other cannot uncoalesce, since they exist in the same locus; a lineage can only uncoalesce with a copied subtree from the parent locus $l$, because a copied subtree represents a 'backbone' for the process rather than a lineage of the new locus. An example of this process is given in Figure S9b.

This process continues backwards in time, until we reach the time of creation of locus $m$.

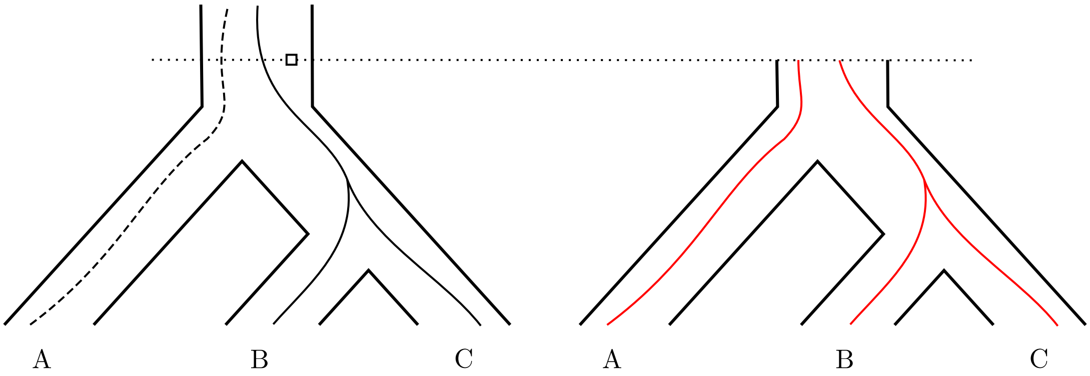


(a) Each lineage begins coalesced with a copied subtree. Here no recombination events occur, so the new haplotype forest is identical to the parent one.

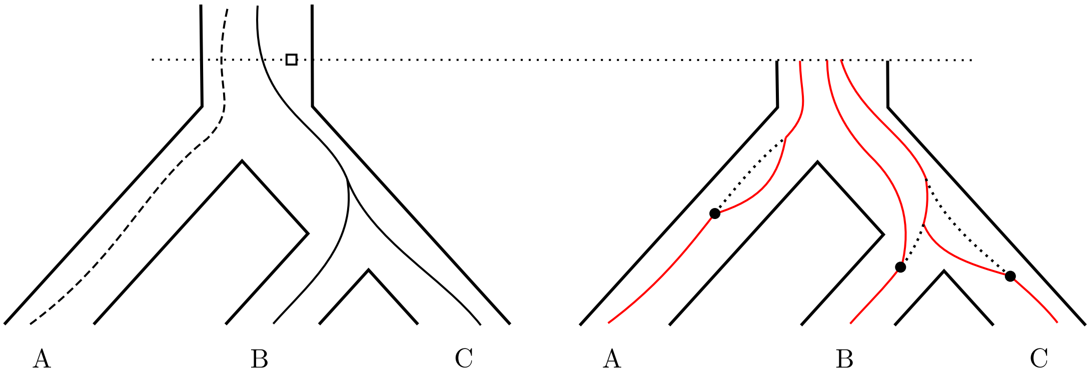


(b) When a recombination event occurs, the lineage 'uncoalesces' from the copied subtree. It may then subsequently coalesce with another lineage or copied subtree.

Figure S9: The linked multispecies coalescent. The subtrees of the haplotype forest in the parent locus are copied to the new locus (dashed black lines). Then, a new multispecies coalescent (in red) is run, with possible recombination events. Finally, one of the red lineages is chosen to be the root of the new haplotype tree. (On the left side of the figure, we use the convention "dashed implies not chosen as haplotype tree".)

This produces the haplotype forest $\mathcal{G}_m$ for the new locus. At this time, we will have a certain number $|\mathcal{G}_m|$ of lineages, some of which may be coalesced with copied subtrees. We choose one uniformly at random, and the descendant tree becomes the new haplotype tree $G_m$.

Observe how this process produces the appropriate results for either extreme of the recombination rate: if $r_r = 0$ (no recombination), then no recombination events occur, and the haplotype tree $G_m$ in the new locus will be a subtree of $\mathcal{G}_l$ (it is possible however that it will not be a subtree of $G_l$, in which case the duplication does not carry a copy of the gene and must be discarded; see the next section). On the other hand, as $r_r \to \infty$ ('infinite' recombination), then any coalescence with $\mathcal{G}_l$ will immediately become uncoalesced, so the haplotype tree $G_m$ will be completely independent from $\mathcal{G}_l$ as desired.

The linked coalescent is very similar to the model of (Slatkin and Pollack, 2006), with a few subtle differences. In our model, we have fixed haplotype information for one locus, and simulate the second locus conditioned on that information, whereas the model of Slatkin and Pollack considers both loci simultaneously. They are also used in different contexts, as (Slatkin and Pollack, 2006) are concerned only with the probability that the two haplotype trees will be identical. Finally, their model is limited to only 3 species; certainly there is no theoretical barrier to extending it to more species, but it quickly becomes too complicated for theoretical analysis, rather than simulation.

This process only produces the haplotype tree and forest in the new locus $m$. The haplotype tree must then be joined back to the haplotype tree in the parent locus, which we detail in the next section.

### 5. Assembling the Full Gene Tree

If a locus generates no duplications or transfers, it will not produce any new loci. Thus almost surely (in the probabilistic sense), the process of generating new unilocus and haplotype trees/forests will stop. When this happens, we assemble the full gene tree by concatenating each haplotype tree in a created locus to the haplotype tree in its parent locus.

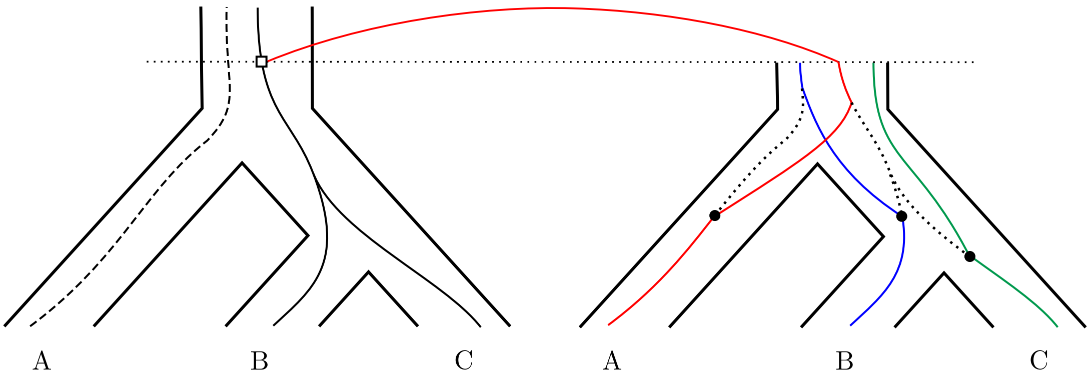


(a) Here the red subtree is chosen to be the new haplotype tree. As it has coalesced with the haplotype tree copied from the parent locus, it is joined directly to that tree at the creation time of the new locus.

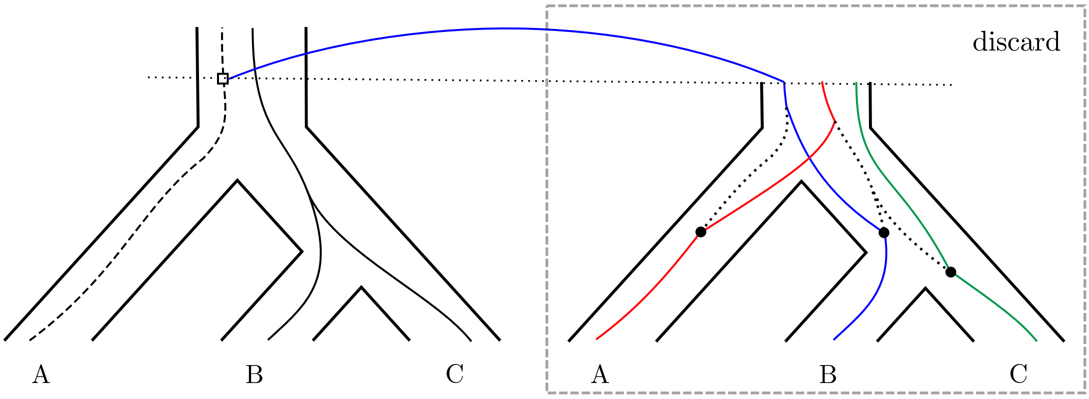


(b) Here the blue subtree is chosen to be the new haplotype tree. It has coalesced with a copied subtree which is not the haplotype tree in the parent locus, so it is discarded.

Figure S10: If a duplication is ancestral, the lineage is joined directly to the subtree with which it has coalesced. (On the left side of the figure, we use the convention "dashed implies not chosen as haplotype tree".)

Recall that linked duplications may be ancestral (the duplicating lineage survives in the parent locus) or non-ancestral (otherwise). Suppose that we wish to re-attach the haplotype tree from a locus $m$ created by a linked duplication, to its parent locus $l$. Once the haplotype tree and forest is generated in locus $m$, we can recognise three cases:

1. The lineage which is chosen to be the haplotype tree $G_m$ in locus $m$ is coalesced (at the time of creation of locus $m$) with (a copy of) the haplotype tree $G_l$ in locus $l$. Then the duplication is an ancestral duplication, and we attach $G_m$ to $G_l$ at the lineage and time where the duplication occurred (Figure S10a, lines 20–21, Algorithm 2).

2. The lineage which is chosen to be the haplotype tree $G_m$ is coalesced with a copied subtree of $\mathcal{G}_l$ which is not the haplotype tree $G_l$. This corresponds to an ancestral duplication of a lineage in locus $l$ which does not carry a copy of the gene. Thus locus $m$ will not carry a copy of the gene, and the locus (along with the duplication, and all loci derived from $m$) is discarded (Figure S10b, lines 22–23, Algorithm 2).

3. The lineage which is chosen to be the haplotype tree $G_m$ is not coalesced with a copied subtree of $\mathcal{G}_l$. Then the duplication is a non-ancestral duplication, and is joined as specified below.

Events which create an unlinked locus (transfers and unlinked duplications) can always be considered to be non-ancestral; since the new locus is independent of the parent locus, the fact that the gene survives in the new locus gives no information about whether it survives in the parent locus (i.e., is ancestral). Therefore the probability of survival in the parent locus is extremely small ($O(\frac{1}{2N})$), and can be safely ignored.

Now, if locus $m$ is created by a non-ancestral event (without loss of generality we will say it is a duplication), the duplicated gene does not survive in the parent locus $l$. Therefore we can treat the duplicating individual as simply another member of the population in the originating branch of $S_l$ at the duplication time $t_m$. We follow this lineage backwards in time (in locus $l$) via the multispecies coalescent to where it coalesces with $\mathcal{G}_l$. Again, there are three cases (lines 24–25, Algorithm 2):

1. The lineage coalesces with the haplotype tree $G_l$. We then attach $G_m$ to $G_l$ at the point where the coalescence occurs (Figure S11a).

2. The lineage coalesces with an element of $\mathcal{G}_l$ which is not the haplotype tree $G_l$. This corresponds to a lineage with no copy of the gene, so locus $m$ is discarded (Figure S11b).

3. The lineage does not coalesce with $\mathcal{G}_l$ by the time of creation of locus $l$. Again, this lineage does not carry a copy of the gene, so locus $m$ is discarded (Figure S11c).

If we do attach $G_m$ to $G_l$, we must finally check that the lineage from the root of $G_m$ to the point where it coalesces with $G_l$ has no losses on it, which we do by running a constant-rate loss process (with rate $r_l$) on that branch. If a loss exists on that branch, the new locus $m$ (and all derived loci) must be discarded (lines 26–28, Algorithm 2).

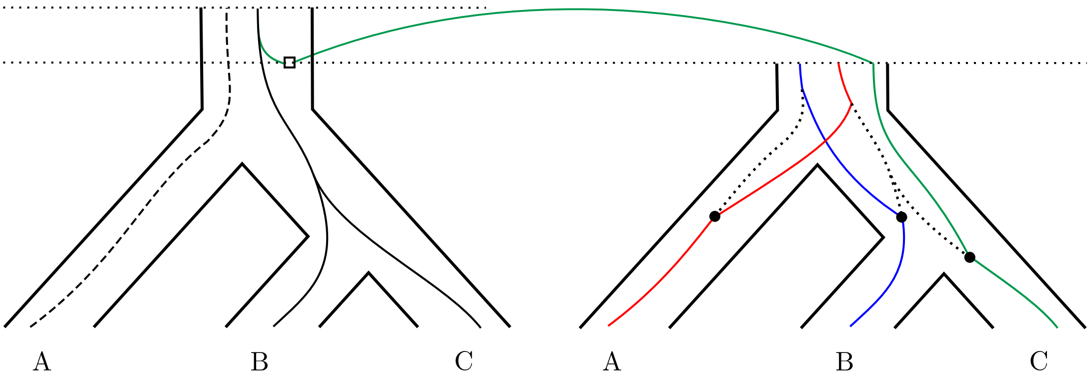


(a) Here the green subtree is chosen to be the new haplotype tree. It coalesces with the haplotype tree in the parent locus, and is joined at the coalescent point.

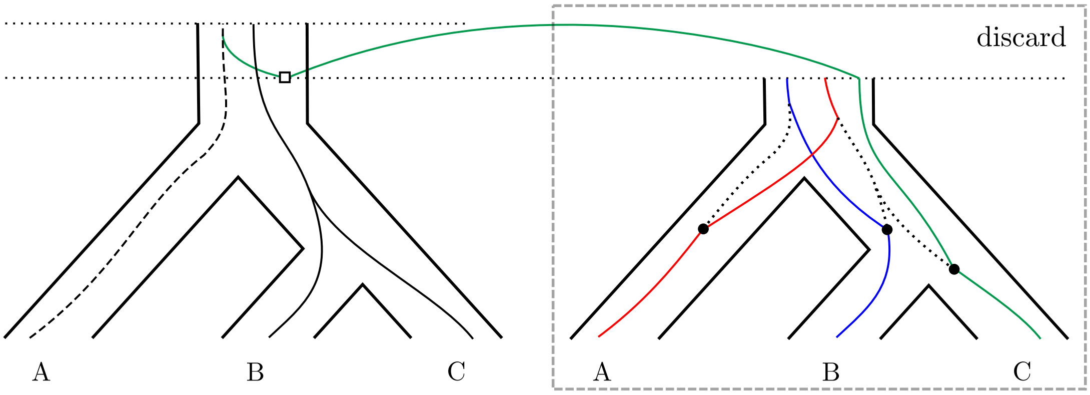


(b) Here the green subtree is chosen to be the new haplotype tree. It coalesces with a lineage which is not in the haplotype tree in the parent locus, so it is discarded.

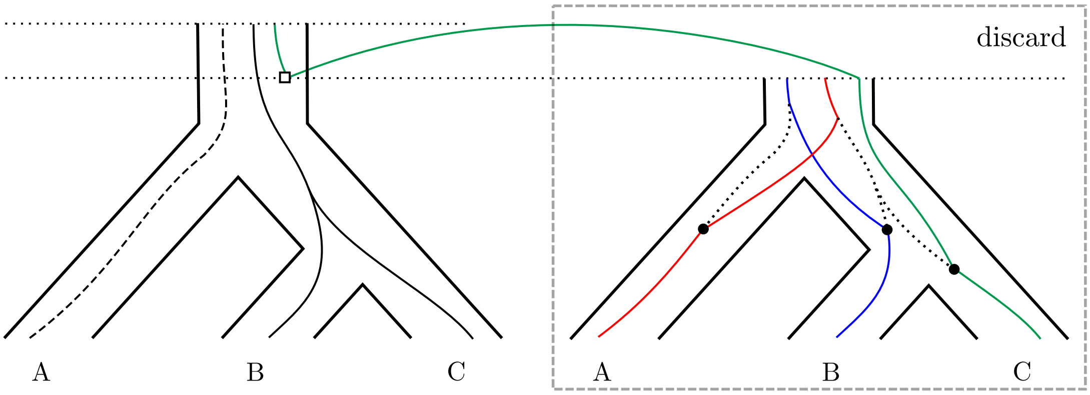


(c) Here the green subtree is chosen to be the new haplotype tree. It does not coalesce by the time of creation of the parent locus, so it is discarded.

Figure S11: If a duplication is non-ancestral, we follow the lineage backwards in time until it coalesces with the haplotype forest in the parent locus. (On the left side of the figure, we use the convention "dashed implies not chosen as haplotype tree".)

This process is done backwards in time order by first attaching (or discarding) each haplotype tree with no child loci to the haplotype trees of their parent loci, and then those to the haplotype trees of their parent loci, and so on until the full gene tree is assembled.

The full pseudocode for this algorithm is given in Algorithms 1 and 2. For clarity, we have chosen to be brief in the specification of the multispecies coalescent processes in lines 14, 18, and 25; for full details, refer to the above discussion. In particular, in line 25, the incomplete coalescent 'joining' may fail to actually join $G_m$ to $G_l^+$, as shown in Figures S11b and S11c, in which case locus $m$ is discarded.

**Algorithm 1** CONSTRUCT__GENE__TREE

**Input**: Species tree $S$ with branch lengths in coalescent units, event parameters $\mathbf{r} = (r_d, r_t, r_l, r_r, p_u)$

**Output**: Gene tree $G$

1: $S_0 \leftarrow S$
2: $G_0 \leftarrow$ multispecies coalescent within $S_0$
3: $\mathcal{G}_0 \leftarrow \{G_0\}$
4: $G_0^+ \leftarrow$ ADD__NEW__LOCI($S_0, G_0, \mathcal{G}_0, G_0, \mathbf{r}$)
5: **return** $G_0^+$

**Algorithm 2** ADD_NEW_LOCI

**Input**: Unilocus tree $S_l$, haplotype tree $G_l$, haplotype forest $\mathcal{G}_l$, partial gene tree $G_l^+$, event parameters $\mathbf{r}$

**Output**: Gene tree $G_l^+$

$\boldsymbol{e^{ud}} \leftarrow$ D coalescent-rate process on $S_l$ with rate parameter $p_u r_d$

$\boldsymbol{e^{t}} \leftarrow$ T coalescent-rate process on $S$ with rate parameter $r_t$, scaled by the proportion of origin species in $S_l$

$\boldsymbol{e^{ld}} \leftarrow$ D coalescent-rate process in the presence of $\mathcal{G}_l$ on $S_l$ with rate parameter $(1 - p_u)r_d$ and recombination rate $r_r$

$\boldsymbol{e^{l}} \leftarrow$ L constant-rate process on $G_l$ with rate parameter $r_l$

**for** $(b, t) \in \boldsymbol{e^{ud}} \cup \boldsymbol{e^{ld}} \cup \boldsymbol{e^{t}} \cup \boldsymbol{e^{l}}$ in time order **do**

 **if** $(b, t) \in \boldsymbol{e^{l}}$ **then**   ▷ here $b \in E(G_l)$

  $g \leftarrow$ a unary node on $G_l^+$ at time $t$ along the branch $b$

  $G_l^+ \leftarrow$ cut $G_l^+$ from $g$

  remove all events from $\boldsymbol{e^{l}}$ below $b$ which occur at a later time

 **else if** $(b, t) \in \boldsymbol{e^{ud}} \cup \boldsymbol{e^{ld}} \cup \boldsymbol{e^{t}}$ **then**   ▷ here $b \in E(S)$

  create a new locus $m$

  $S_m \leftarrow$ subtree of $S_l$ rooted at $b$ at time $t$

  **if** $(b, t) \in \boldsymbol{e^{ud}} \cup \boldsymbol{e^{t}}$ **then**

   $\mathcal{G}_m \leftarrow$ incomplete coalescent within $S_m$ to time $t$

   $G_m \leftarrow$ an element of $\mathcal{G}_m$

  **else if** $(b, t) \in \boldsymbol{e^{ld}}$ **then**

   $\mathcal{G}'_m \leftarrow$ all subtrees of $\mathcal{G}_l$ within $S_m$

   $\mathcal{G}_m \leftarrow$ linked coalescent in the presence of $\mathcal{G}'_m$ within $S_m$ to time $t$, with recombination rate $r_r$

   $G_m \leftarrow$ an element of $\mathcal{G}_m$

  **if** $(b, t) \in \boldsymbol{e^{ld}}$ and $G_m$ has coalesced with a subtree of $G_l$ at time $t$ **then**

   attach $G_m$ to $G_l^+$ at time $t$

  **else if** $(b, t) \in \boldsymbol{e^{ld}}$ and $G_m$ has coalesced with a subtree of $\mathcal{G}_l$ at time $t$ **then**

   discard locus $m$

  **else**

   $G_l^+ \leftarrow$ incomplete coalescent 'joining' of $G_m$ to $G_l^+$ in the presence of $\mathcal{G}_l$ within $S_l$ (keep $G_l^+$)

   $\boldsymbol{e^{l}} \leftarrow$ L constant-rate process from root of $G_m$ to $G_l^+$ with rate parameter $r_l$

   **if** $|\boldsymbol{e^{l}}| > 0$ **then**

    discard locus $m$

  $G_l^+ \leftarrow$ ADD_NEW_LOCI$(S_m, G_m, \mathcal{G}_m, G_l^+, \mathbf{r})$

**return** $G_l^+$
